# Brain hierarchy score: Which deep neural networks are hierarchically brain-like?

**DOI:** 10.1101/2020.07.22.216713

**Authors:** Soma Nonaka, Kei Majima, Shuntaro C. Aoki, Yukiyasu Kamitani

**Affiliations:** Faculty of Integrated Human Studies, Kyoto University, Sakyo-ku, Kyoto 606-8501, Japan; Graduate School of Informatics, Kyoto University, Sakyo-ku, Kyoto 606-8501, Japan; ATR Computational Neuroscience Laboratories, Kyoto 619-0288, Japan

**Keywords:** deep neural network, visual cortex, functional magnetic resonance imaging (fMRI), neural decoding, visual hierarchy

## Abstract

Achievement of human-level image recognition by deep neural networks (DNNs) has spurred interest in whether and how DNNs are brain-like. Both DNNs and the visual cortex perform hierarchical processing, and correspondence has been shown between hierarchical visual areas and DNN layers in representing visual features. Here, we propose the brain hierarchy (BH) score as a metric to quantify the degree of hierarchical correspondence based on the decoding of individual DNN unit activations from human brain activity. We find that BH scores for 29 pretrained DNNs with varying architectures are negatively correlated with image recognition performance, indicating that recently developed high-performance DNNs are not necessarily brain-like. Experimental manipulations of DNN models suggest that relatively simple feedforward architecture with broad spatial integration is critical to brain-like hierarchy. Our method provides new ways for designing DNNs and understanding the brain in consideration of their representational homology.

## Introduction

The design of deep neural networks (DNNs) is typically based on brain-like multi-stage hierarchical structures. DNNs have been reported to achieve human-level performance in some cognitive tasks, including image recognition. The structural and behavioral similarities between DNNs and biological brains have triggered investigation on how DNNs and biological brains are “functionally” similar. A large number of studies have suggested that task-optimized DNNs acquire similar representations to task-related brain regions. For example, neuronal responses in the monkey inferior temporal (IT) cortex, which underlies object recognition, are predicted accurately by unit activation patterns of DNNs trained to perform visual tasks (Cadieu et al., 2014; Khaligh-Razavi and Kriegeskorte, 2014; Yamins et al., 2014). DNN’s activations can also successfully explain human functional magnetic resonance imaging (fMRI) responses (Eickenberg et al., 2017; Guclu and van Gerven, 2015; Khaligh-Razavi and Kriegeskorte, 2014). Moreover, recent studies have demonstrated hierarchical correspondence of representations, or hierarchical homology, between DNNs and biological brains (Cichy et al., 2016; Guclu and van Gerven, 2015; Horikawa and Kamitani, 2017). Horikawa and Kamitani (2017) reported that human fMRI responses to visual images can be decoded (translated) into unit activations of a DNN (AlexNet) responding to the same images. The brain areas that best predict unit activations in a DNN layer are reported to gradually shifted from lower (e.g., V1, V2, and V3) to higher visual areas (e.g., V4, lateral occipital complex, fusiform face area, and parahippocampal place area) as the target DNN layer shifts from lower to higher. This finding indicates functional similarity of the hierarchical representations between DNNs and biological brains. Why is the functional similarity between DNNs and the brain important? Brain-like hierarchical representations have the potential to realize DNNs that exhibit more human-like behavior, and to overcome the limitations of the current task-optimized DNNs. To design and train DNNs to develop hierarchical representations similar to the human brain, DNNs are expected to share the feature spaces with the human brain and emulate its internal information processing. Such DNNs would behave more similarly to humans than DNNs trained to maximize the task performance. Potential advantages of the brain-like DNNs include robustness against adversarial attack (Szegedy et al., 2013), generalizability across datasets with various types of image distortion (Geirhos et al., 2018), and realistic behavior patterns as surrogates of humans. Moreover, brain-like representations and hierarchy are considered to be helpful for the development of artificial general intelligence. In addition to the potential benefits of the development of DNNs, brain– DNN functional similarity measurement may advance our understanding of the brain by providing better computational models (Yamins and DiCarlo, 2016) or experimental tools generating optimal stimuli for neurons (Bashivan et al., 2018; Ponce et al., 2019). Thus, quantitative measures for evaluating how a given DNN has similar representations to the brain and how the representations develop brain-like hierarchies may have a range of valuable applications.

In a previous study (Schrimpf et al., 2018) the Brain-Score was proposed as a quantitative metric of the brain–DNN similarity of representations. The Brain-Score evaluates how accurately neuronal responses in primate visual areas were predicted from DNN unit activation patterns. This previous study systematically compared the similarities between various DNNs for image recognition and representations in middle and higher visual areas (V4 and IT cortex), reporting a positive correlation between object classification accuracy on the ImageNet dataset (i.e., ImageNet top-1 accuracy) and Brain-Score across 69 DNNs. This correlation, however, was weaker for more recently developed high-performance DNNs (DNNs with ≥ 70% accuracy), suggesting that performance improvement does not necessarily lead to brain-like DNNs. Although the Brain-Score captures the similarity of representations between DNNs and individual brain areas, it does not evaluate hierarchical homology across layers/brain areas between DNNs and the brain. Since multistage hierarchical processing is considered to play a vital role in perception and cognition, quantitative evaluation of the brain–DNN hierarchical homology may provide a sensitive comparison of functional similarity between DNNs and the human brain.

In the current study, to quantitatively evaluate the degree of brain–DNN hierarchical homology across DNN layers and cortical areas, we propose an index called the brain hierarchy (BH) score. The BH score is designed to capture the extent to which DNN layers of a given DNN are aligned with the hierarchy of the brain. To compute the BH score, individual DNN units are characterized by decoding of DNN unit activation from brain activity (Horikawa and Kamitani, 2017); we predict the activation of each unit from fMRI responses in one of five brain areas (regions of interest; ROIs) covering the entire cortical hierarchy (V1, V2, V3, V4, and higher visual cortex [HVC]), then identify the ROI showing the highest decoding accuracy among the five ROIs (hereafter referred to as the “top ROI”; Figure 1A). The BH score is defined as the Spearman rank correlation coefficient between the layer number and the top ROI across units in the given DNN (Figure 1B). When the top ROIs of DNN units monotonically increase with respect to their layer number in a given DNN, the DNN shows the highest BH score. As a complementary measure, by exchanging DNN layers and brain areas, the BH score is also calculated based on encoding models that predict fMRI voxel values from DNN unit activities to examine the robustness of the results.

**Figure 1.**
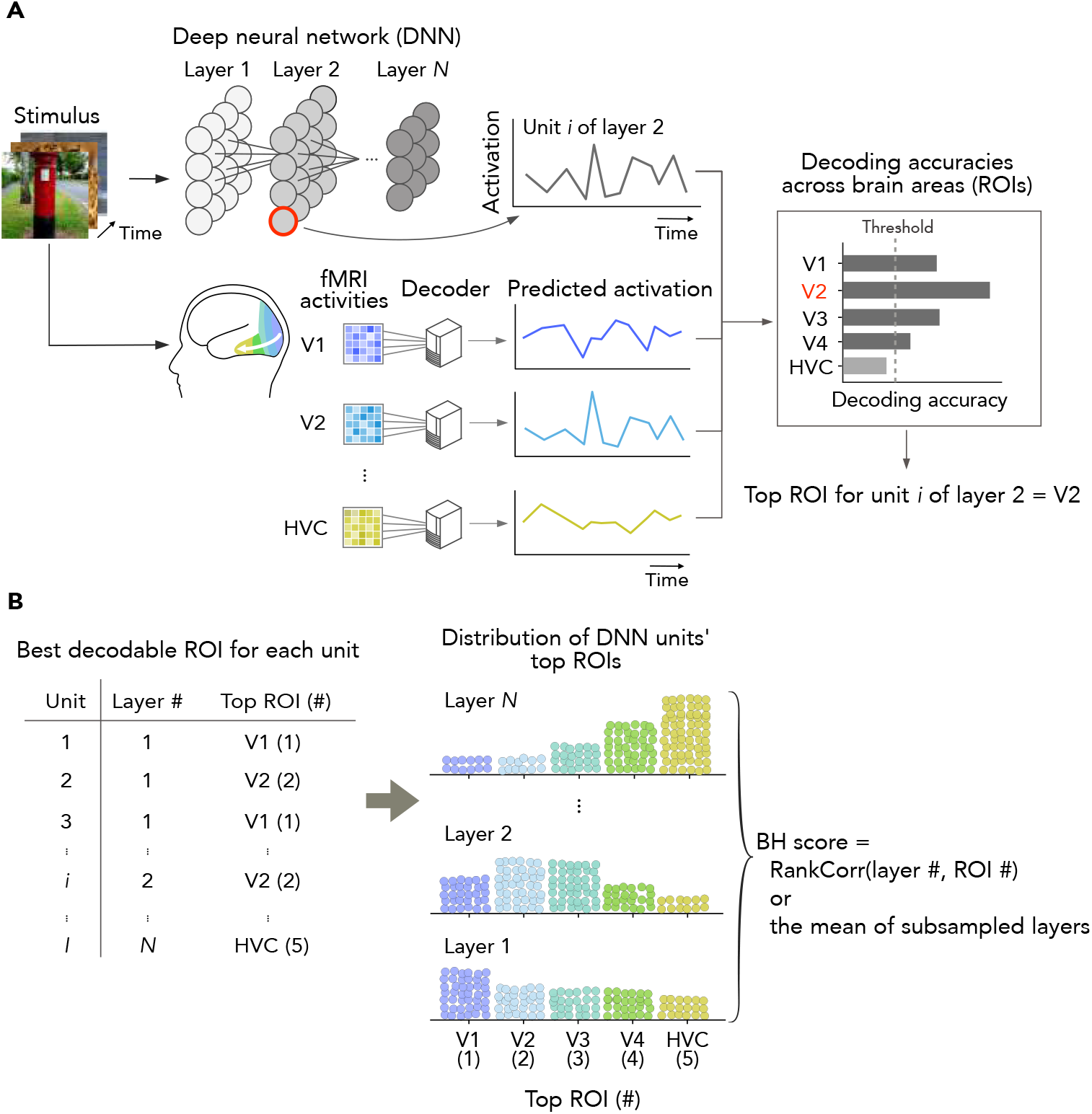
Evaluation of hierarchical homology. **(A)** Identification of the best decodable ROI (top ROI) for each DNN unit. To characterize individual deep neural network (DNN) units using human functional magnetic resonance imaging (fMRI), the responses of each DNN unit given natural images were predicted (decoded) from the fMRI responses to the same images in each of five ROI in the visual cortex. The ROI showing the best decoding accuracy was assigned as the top ROI for the unit. The units that were not predicted by any of the ROIs above a statistical threshold were excluded from further analyses. **(B)** Top ROI distributions for DNN layers and the brain hierarchy (BH) score. The distributions of the top ROIs for individual units are visualized for each DNN layer. The BH score was evaluated using the Spearman rank correlation between the layer and the ROI numbers. When DNNs to be compared have different number of layers, we randomly subsampled five layers, and calculated the mean BH score across random selections.

Using the distributions of the top ROIs and the BH scores, the degree of hierarchical similarity to the brain is examined in 29 representative DNNs pre-trained on an object classification task, including AlexNet, the VGG family, the ResNet family, the DenseNet family, and the Inception family (see Transparent Methods: “Deep neural networks” and Table 1). DNNs with the same architectures and random weights are also studied to examine the effects of training. We investigate the correlations between BH scores and ImageNet top-1 accuracy to determine whether high-performance DNNs are hierarchically more brain-like, while testing the robustness of the BH score and the relationship with other measures of brain–DNN similarity.

**Table 1.**
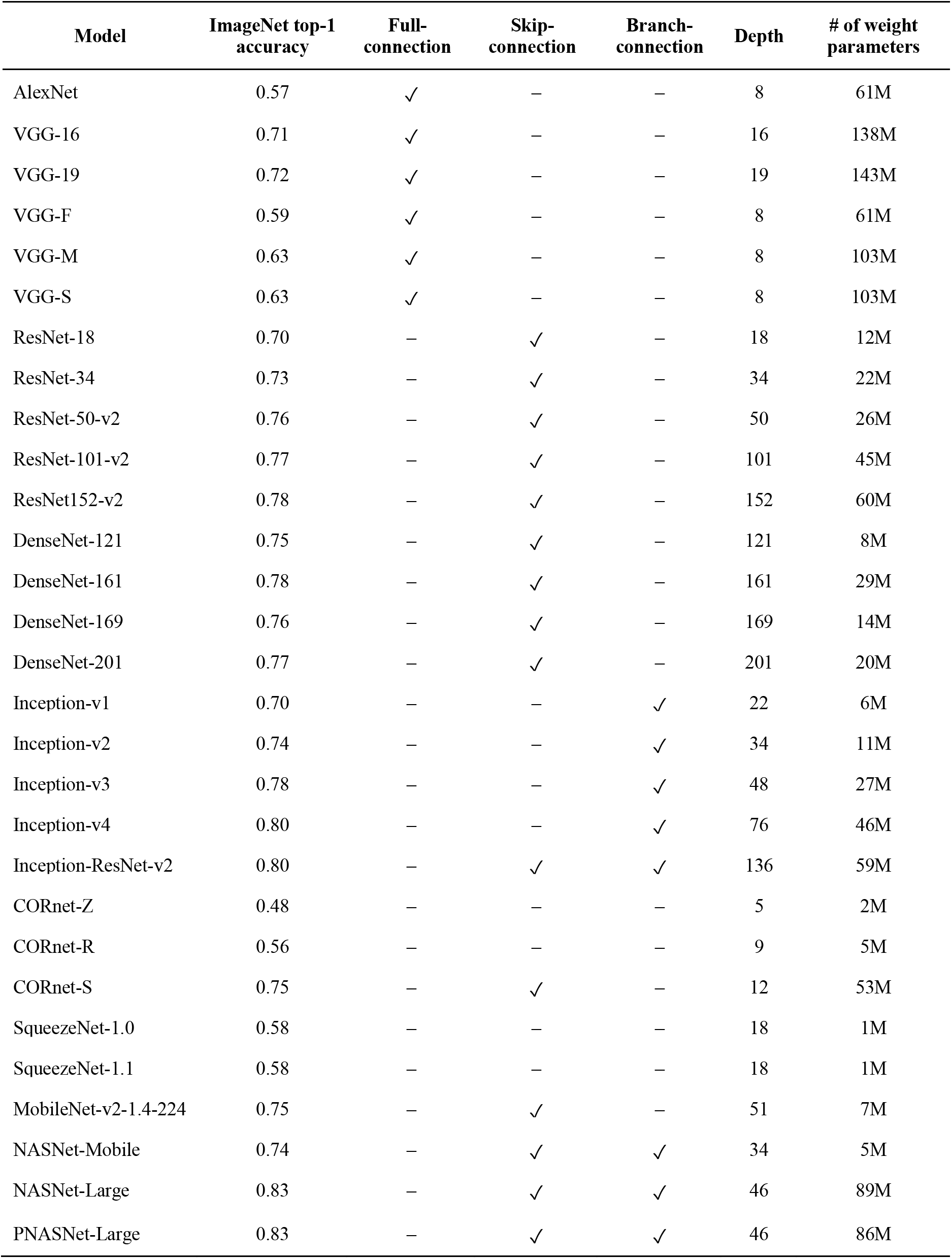
29 deep neural networks compared in the current study.

We then seek to identify architectural characteristics of DNNs associated with the degree of hierarchical homology. We focus on five representative components of DNN architecture: the presence of fully-connected (FC) layers, the presence of branch-connections, the presence of skip-connections, the number of DNN layers, and the number of weight parameters. These architectural components are compared with BH scores among the 29 pre-trained DNNs. Then, some of the components are experimentally manipulated to evaluate the effects on BH scores in trained DNNs with otherwise identical architectures. The code and the fMRI data to compute the BH score for novel DNNs are provided for public use at repositories (see Transparent Methods: “Data and Software Availability”).

## Results

### Brain hierarchy score

The brain hierarchy (BH) score is designed to evaluate a deep neural network (DNN) for object recognition in terms of the hierarchical similarity to the human brain by determining the correspondence of each unit in each DNN layer to one of the hierarchically ordered visual cortical areas (regions of interests, ROIs). To consider hierarchical representations of DNNs, we included representative layers of each DNN in the analysis: the first layer, the last fully-connected (FC) layer (referred to as the “category layer” in the current study), the other FC layers, convolutional layers that do not belong to submodules, and the output layers of submodules (i.e., convolutional-, skip- or branch-block). Hereafter, “layer” refers to the representative layer, unless otherwise stated. The layers in each DNN are numbered from input to output. Each DNN unit is labeled by the layer number it belongs to. The brain ROIs include V1, V2, V3, V4, and a combined region of ventral object-responsive areas (higher visual cortex [HVC]; see Transparent Methods: “Region of interests”) and each is assigned an ROI number: V1 (1), V2 (2), V3(3), V4 (4), and HVC (5).

For each DNN unit, we identified the ROI with the best linear decodability (i.e., the ROI whose functional magnetic resonance imaging (fMRI) voxel pattern can best predict the DNN unit activity using a linear decoder given the same input image) which we refer to as the “top ROI.” DNN units that were poorly predicted from any of the ROIs were excluded in later analyses to improve sensitivity of the BH score to the hierarchical representations (Figure 1A; see Transparent Methods: “Brain hierarchy score” for details). For the first DNN layer in which linear spatial filtering is performed, the absolute values of the raw unit activations were used as targets for decoding, because fMRI signals are known to be sensitive to deviations from baseline luminance but not luminance itself, even at V1, presumably reflecting complex cell-like early nonlinear processing (Haynes et al., 2004; see Transparent Methods: “Decoding analysis”). We used the unit selection and the first layer nonlinearity as default settings. We discuss how these settings affected BH scores in a later section. Repeating the procedures for the DNN units used in the decoding analysis, we obtained the distributions of the top ROIs for each layer for visualization (Figure 1B). If a DNN has hierarchical representations that are similar to those of the brain, the distribution of top ROIs should gradually shift from lower to higher cortical areas as the DNN layer increases.

The BH score quantifies this gradual shift of top ROIs, exhibiting a high value if the distribution monotonically shifts with the layer number, and a low value if 1) the peak does not monotonically shift or 2) the distribution is flat or multimodal with highly variable top ROIs at each layer. Because the number of layers differs across DNNs and brain ROIs could be delineated differently, a linear relationship would not be expected between the layer and ROI numbers. Thus, we used the Spearman rank correlation between the layer and ROI numbers, measuring the degree of the monotonic relationship between the two variables.

The BH score is defined as the Spearman rank correlation coefficient between the layer number and the top ROI number across DNN units. To compute the Spearman rank correlation coefficient, the values of input variables are transformed into fractional ranks. Here, we denote the layer number and top ROI number of the *i*-th DNN unit (*i* = 1,*I*,) by 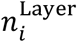 and 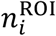, respectively. The fractional ranks of the layer number and the top ROI number of the *i*-th DNN unit are respectively defined as

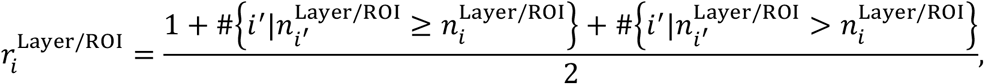

where #{·} denotes the number of the members in the set {·}. The BH score is then computed as

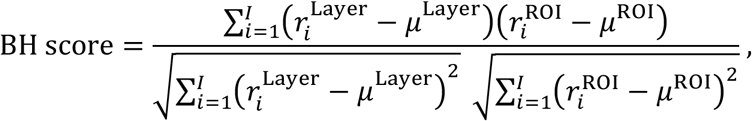

where *μ*^Layer^ and *μ*^ROI^ are the means of 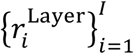 and 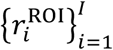, respectively. Because this measure is based on the correlation calculated with individual units, not just on the peaks of the distributions, the variability in the distribution is taken into account, so that highly fluctuating top ROIs in each layer lead to low BH scores. Randomly selected 1000 units per layer were used in the decoding analysis. The remaining DNN units after excluding poorly predicted ones were used. To match the number of layers used to compute the BH score across DNNs, the BH score was computed with the first, last, and randomly selected three intermediate layers. The random layer selection and BH score computation were repeated 100 times, and the mean score is reported.

Our proposed method is based on the decoding of individual DNN units, enabling a DNN to be comprehensively characterized at the finest level. We assume that DNN units serve as proxies for feature detecting neurons that are distributed in the visual cortex. Whereas fMRI voxels lack the resolution to isolate a single neuron, the distributed representation of the same feature could be read out by combining many voxels with optimal weights (“ensemble feature selectivity”; Kamitani and Tong, 2005). A complimentary evaluation can be performed with the encoding analysis by exchanging fMRI voxels and DNN units. Although we mainly used the decoding-based BH score, some results are shown with both methods to examine robustness of the BH score.

In the current study, we performed decoding analysis using fMRI data collected by Shen et al. (2019) (see Transparent Methods: “fMRI dataset”). This dataset is composed of fMRI activity of three subjects viewing 1250 natural object images from ImageNet (2011, fall release; Deng et al., 2009). fMRI responses to 1200 and 50 natural object images were separately used as training and test data for the decoders. The top ROIs were defined using fMRI data from individual subjects separately. Thus, the BH score could be computed using the fMRI data from each subject. We report the consistency of BH scores across the three subjects. Unless stated otherwise, we calculated the Spearman rank correlation coefficient by pooling all top ROIs across three subjects.

### Comparison of BH scores across DNNs

We compared BH scores between 29 representative DNNs (Table 1). All DNNs were pre-trained to classify object categories on ImageNet (Deng et al., 2009). These DNNs show different tendencies in their distribution of top ROIs of DNN layers (Figure 2; also see Figure S1). For example, DNNs with FC layers (e.g., AlexNet; Krizhevsky et al., 2012) and the VGG family (Figure S1) show a clear shift of top ROI distributions from lower to higher visual areas along with the hierarchy of DNN layers. Unit activations in early convolutional layers were better predicted from the lower visual areas, those in late convolutional layers were better predicted from V3, and those in the FC layers were better predicted from V4 or HVC. This clear hierarchical correspondence between the DNN layers and the visual cortical areas leads to high BH scores (e.g., BH score = 0.50 for AlexNet). Inception-v1 (Szegedy et al., 2014), which consists of blocks with branch-connections (inception module), shows flat distributions of top ROIs and had a low BH score (BH score = 0.27). Such flat distributions are also observed in other DNNs with branch-connections (e.g., the Inception family, NASNet [Zoph et al., 2018] and PNASNet [Liu et al., 2018]; Figure S1). One possible reason for the flattened distributions of top ROIs is that different sizes of convolution and pooling operations are parallelly applied in branch-connections and DNN units corresponding to different brain areas tend to co-exist within the individual layers. In DNNs with skip-connections (residual module) such as ResNet-152-v2 (He et al., 2015), NASNet-Large (Zoph et al., 2018) and variants of these DNNs (e.g., the DenseNet family and PNASNet-Large [Liu et al., 2018]), the peaks of top ROI distributions non-monotonically swing between V1 and V3, leading to low BH scores (e.g., 0.14 for Resnet-152-v2 and 0.07 for NASNet-Large; Figure S1). This tendency may be caused by bypassing of representations via skip-connections: representations similar to lower visual areas are bypassed from low to higher DNN layers by the skip-connections, and the higher layers tend to include units that are well predicted from fMRI responses in the lower visual areas.

**Figure 2.**
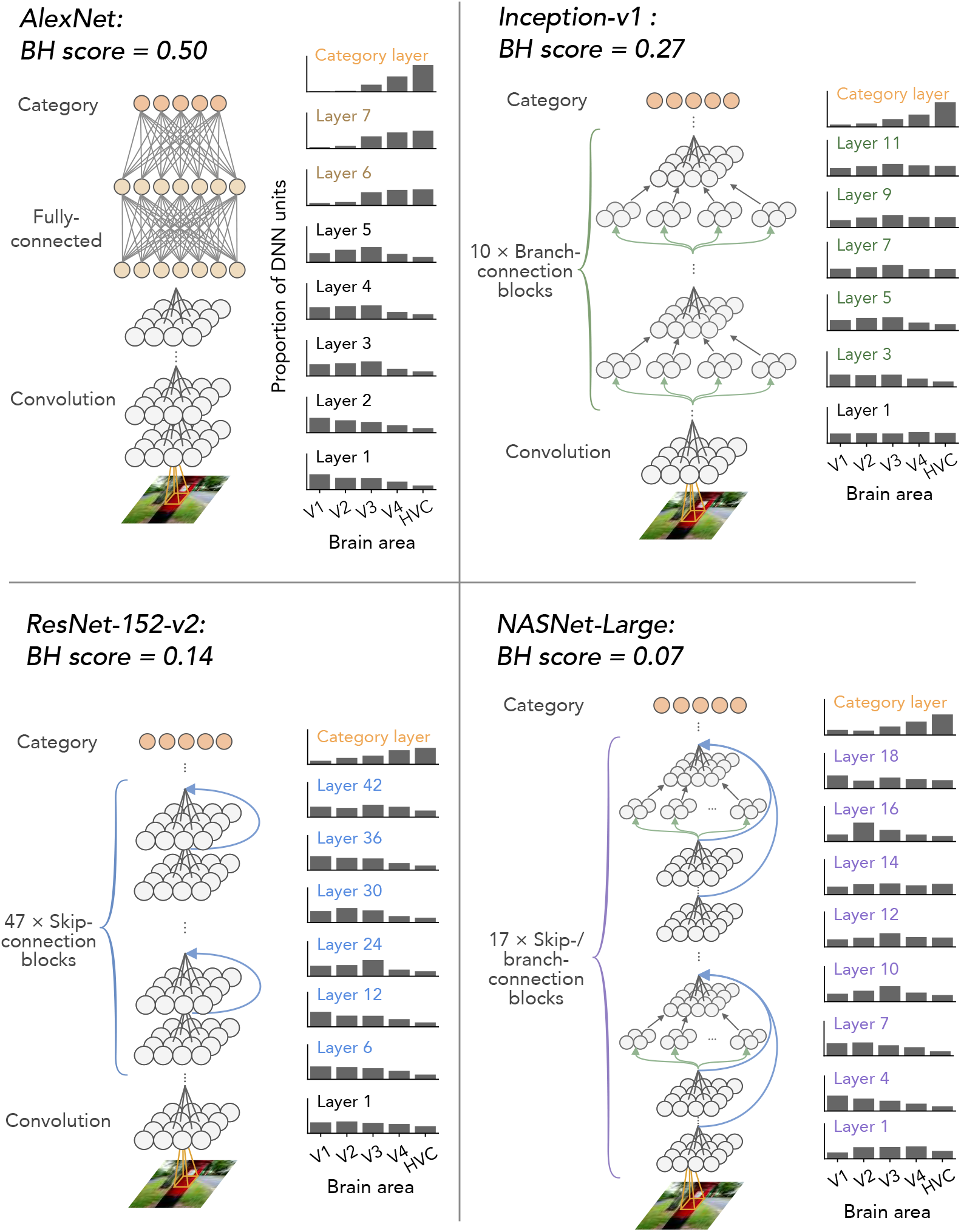
Top ROI distributions and BH scores for representative DNNs. The distributions of top ROIs for each layer of AlexNet, Inception-v1, ResNet-152-v2, and NASNet-Large are shown with schematics of their architectures. Histograms are normalized for each layer by the total number of counted DNN units. To match the number of layers used to calculate the BH score across DNNs, BH scores were computed by randomly choosing five DNN layers. The mean BH scores across 100 random selections are shown above.

By comparing these 29 DNNs, we found that DNNs with simple architecture (e.g., AlexNet, the VGG family, and the CORnet family) exhibit relatively high BH scores (Figure 3). In contrast, DNNs with elaborate architecture and high image recognition performance, such as the DenseNet family, the ResNet family, and the Inception family, show low BH scores. BH scores are negatively correlated with the ImageNet top-1 accuracies across the DNNs (Figure 3; *ρ* = −0.73, permutation test, *p* < 0.01). Thus, BH scores show an opposite tendency to Brain-Score, indicating that high-performance DNNs are *not* brain-like if hierarchy is considered.

**Figure 3.**
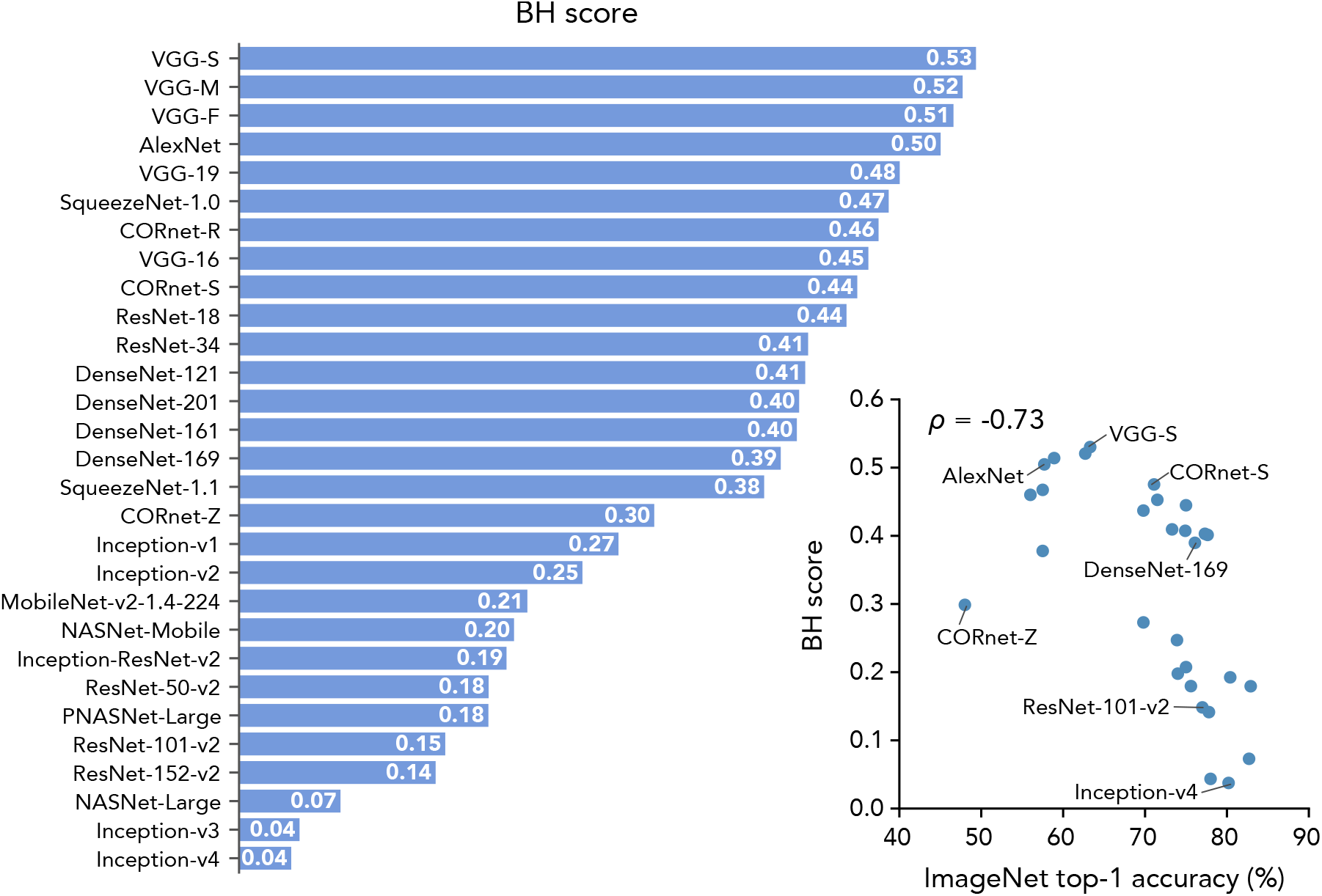
BH scores and ImageNet top-1 accuracies. The BH scores for the 29 DNNs are shown in descending order of the BH score. In the right panel, the BH scores for the 29 DNNs are plotted against ImageNet top-1 accuracies.

To examine whether this negative correlation is robust to the measurement of image recognition performance, we evaluated DNNs’ image recognition accuracies on image datasets other than ImageNet: Caltech-101 (Fei-Fei et al., 2007) and Caltech-256 (Griffin et al., 2007). A multinomial logistic regression classifier was trained using the second last layer of each DNN as input. The classifiers were trained and tested with the training and test data of each dataset. BH scores are also negatively correlated with image recognition accuracy evaluated on Caltech-101 data and that evaluated on Caltech-256 data (Figure S2).

To examine whether training is necessary for DNNs to yield brain-like hierarchical representations, we compared the degree of hierarchical homology between trained and untrained (i.e., with random weights) versions of the 29 DNNs. To construct untrained DNNs, we used randomly initialized weights provided in the original implementation of the DNNs. Overall, the untrained DNNs exhibit lower BH scores than DNNs trained on ImageNet (Figure 4). The monotonic shift of the distribution of the top ROIs along the hierarchy of DNN layers was deteriorated by randomization of DNNs’ weights (Figure 4A). The unit activations in the untrained DNNs are less predictable, particularly in higher layers (e.g., layers 6 and 7, and the category layer in AlexNet), from the fMRI responses in higher visual areas (V4 and HVC) compared with those in the trained DNNs. This effect resulted in flattened, middle ROI-peaked, or lower ROI-peaked distributions of the top ROIs in each layer, and degraded the BH scores of DNNs with random weights. Similar degradation of BH scores was observed in the majority of tested DNNs (Figure 4B). The results indicate that the hierarchical homology of DNNs to the brain does not arise only from their architectures, but also requires training with image data.

**Figure 4.**
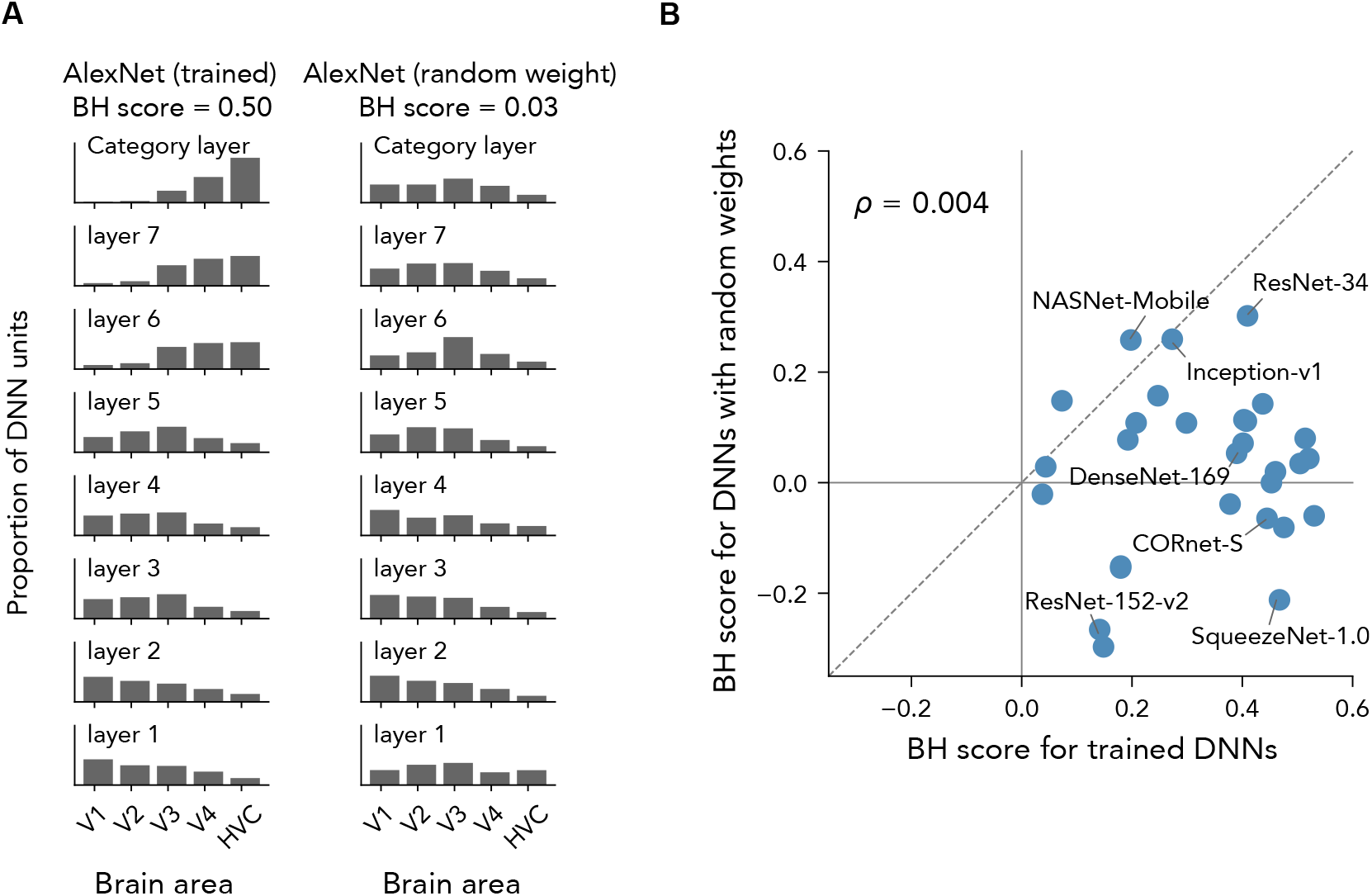
BH scores for trained and untrained DNNs. **(A)** The top ROI distributions and BH scores for the original trained AlexNet and the untrained AlexNet. Both DNNs had the same architecture but the untrained AlexNet had random weights. **(B)** BH scores for trained and untrained DNNs. The BH scores for the original trained 29 DNNs and for the untrained DNNs are shown in a scatter plot.

### Robustness of the BH score

We defined the BH score based on several optional choices of procedures. Here we consider the robustness of the BH score to some of these choices. In the calculation of the BH score, we excluded DNN units whose decoding accuracies did not exceed a predefined threshold for any ROIs (student’s *t*-test, *p* < 0.05). On average, 42.9% of DNN units were excluded by this procedure. We computed the BH score without this unit exclusion and compared it with the original BH score (Figure S3A). BH scores with and without unit selection were highly correlated across the 29 DNNs (*ρ* = 0.88). In addition, we found highly correlated scores when raw unit activations instead of absolute values were used for the first layer (Figure S3B, *ρ* = 0.98). While the scores with and without these procedures were strongly correlated, the original BH scores tended to take a wide range of values, indicating that these procedures are effective at detecting monotonic shifts with more sensitivity.

We also examined the consistency of BH scores across fMRI data from individual subjects. We computed BH scores using fMRI data for each subject, and similar scores were observed consistently across subjects (Figure S4). The results suggest that BH scores do not depend on the particular brain used for calculation.

We further examined whether similar comparison results were consistently observed with encoding analysis. Here, we performed encoding analysis in which individual voxels in each ROI were predicted from each DNN layer (see Transparent Methods: “Encoding analysis”), and quantified the degree of brain–DNN hierarchical homology based on encoding accuracies using the same procedure as that for the BH score (Figures S5A and S6). The BH score based on decoding analysis is positively correlated with the BH score based on encoding analysis (Figure S5B; *ρ* = 0.55), indicating robustness to the choice of the analysis method.

### Similarity between DNNs and single brain regions

Whereas the BH score was negatively correlated with image recognition performance across DNNs, previous studies reported a positive correlation between their proposed similarity measure and image recognition performance (Schrimpf et al., 2018; Yamins et al., 2014; Yamins and DiCarlo, 2016). Because these previous studies evaluated the similarity of representations between DNNs and specific brain areas, such as V4 and IT, whether hierarchy is considered or not may constitute a critical difference. Another possible source of this discrepancy is the difference in the direction of prediction, because these previous studies defined similarity as the encoding accuracy for predicting neuronal responses from DNN unit activity patterns. However, it should be noted that similar comparison results were observed between decoding and encoding analysis, as shown above.

The discrepancy may have also arisen from differences in measurement modality, because previous studies used electrophysiological recordings of neuronal responses from monkey cortex. To examine whether the modality is critical, we performed similar encoding analysis with our fMRI data. In accord with the brain regions examined in previous studies (Schrimpf et al., 2018; Yamins et al., 2014; Yamins and DiCarlo, 2016), we selected V4 and HVC for this encoding analysis. The responses of individual voxels in V4 and HVC were predicted from the activations of DNN units in each DNN (Figure 5; Transparent Methods: “Encoding analysis”). Consistent with the previous finding, we observed a positive correlation between ImageNet top-1 accuracy and encoding accuracy across DNNs (*ρ* = 0.76 for V4; *ρ* = 0.68 for HVC). Thus, the results are robust to differences in measurement modality, suggesting that the inclusion of hierarchy is critical for similarity evaluation.

**Figure 5.**
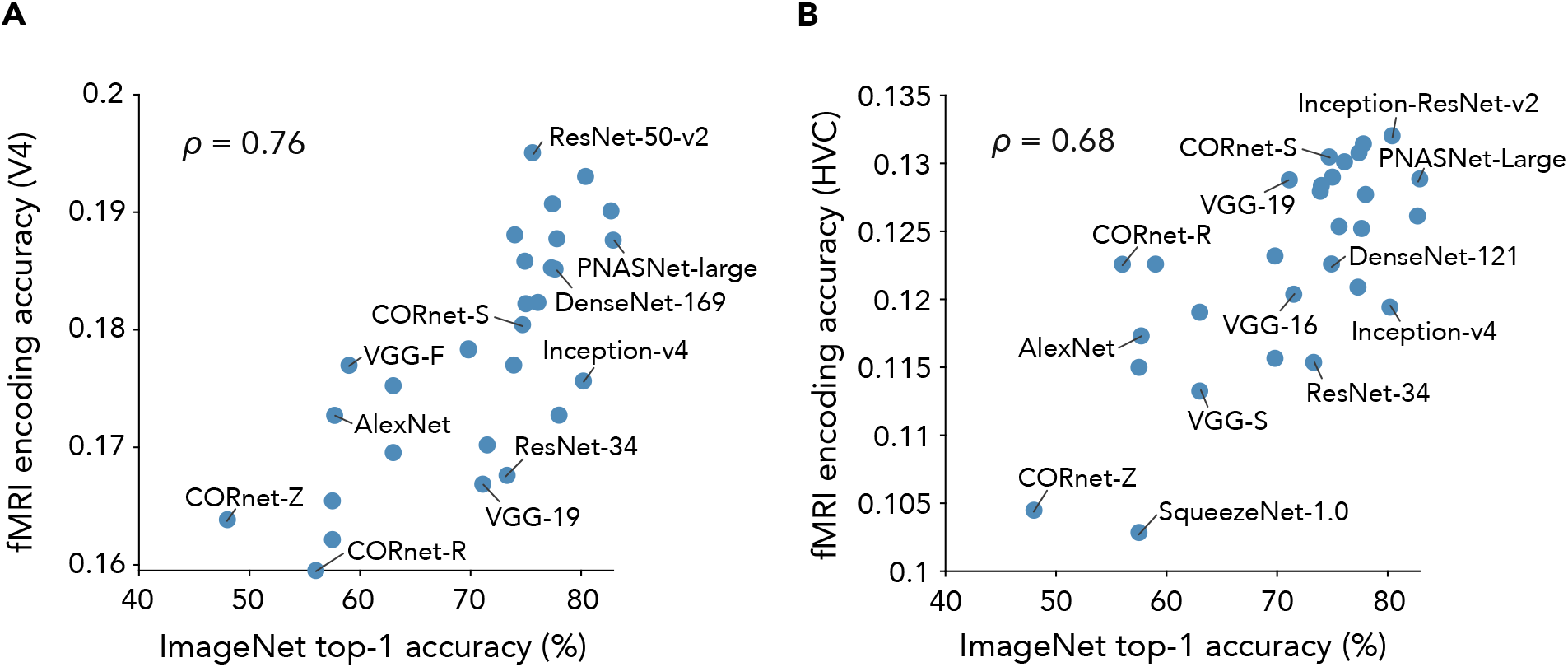
fMRI encoding accuracy and ImageNet top-1 accuracy. **(A)** fMRI V4 encoding accuracy and ImageNet top-1 accuracy. Responses of Individual voxels in V4 were predicted from unit activations of each DNN. The mean prediction accuracies (encoding accuracies) across the voxels are plotted against ImageNet top-1 accuracies. **(B)** fMRI higher visual cortex (HVC) encoding accuracy and ImageNet top-1 accuracy. The mean encoding accuracies for HVC are plotted against ImageNet top-1 accuracies.

Recent studies have used representational similarity analysis to evaluate the similarity between a DNN and human IT using fMRI, reporting a negative correlation between the representational similarity to human IT and ImageNet top-1 accuracy across DNNs (*r* = −0.47 in Jozwik et al. [2019] and *r* = −0.38 in Storrs et al. [2020]). We replicated the results using fMRI data in the current study (*r* = −0.55; Figure S7). Although these results are based on individual brain areas, not hierarchy, the lower similarity of high-performance DNNs to the higher visual areas (human IT or HVC) might contribute to poor hierarchical correspondence, accounting for the negative correlation between BH scores and image recognition performance. However, it remains unclear why representational similarity and predictive (encoding or decoding) accuracy exhibited opposite correlations to image recognition performance when measured in individual areas.

### BH scores and DNN architectural components

What components of architecture are critical for brain-like hierarchical representations in DNNs? We examined how BH scores are explained by five representative components of DNN architectures: the presence of FC layers, the presence of skip-connections, the presence of branch-connections, the total number of convolutional and FC layers (i.e., depth), and the number of weight parameters. Note that to define the depth of a DNN, we counted not only the representative layers but also those not used in the computation of the BH score. For the first three components, we compared the mean BH score between the presence and the absence of each component. Correlations with the BH score were calculated for the depth and weight parameters. Because only two DNN (CORnet-R and CORnet-S) had recurrent connections among the DNNs we tested in the current study, the difference due to the presence of recurrent connections was not quantitatively examined (BH scores for CORnet-R and CORnet-S are 0.44 and 0.41, which is relatively high among the tested DNNs).

DNNs with FC layers exhibit markedly higher BH scores than those without FC layers (*d’* = 1.29 [*d*-prime: the mean difference normalized by the standard deviation]; Figure 6A), while those with skip-connections and branch-connections show moderately lower BH scores: *d’* = −0.30 for skip-connections and *d’* = −0.65 for branch-connections (Figure 6A). We also found a negative correlation between BH scores and depth (*ρ* = −0.63; Figure 6B), and a weak positive correlation between BH scores and the number of weight parameters (*ρ* = 0.15; Figure 6B). Complementary regression analysis, in which BH scores were modeled as a linear weighted sum of variables parameterizing the five components, also indicated a greater contribution of the presence of FC layers compared with the other components (Figure 6C).

**Figure 6.**
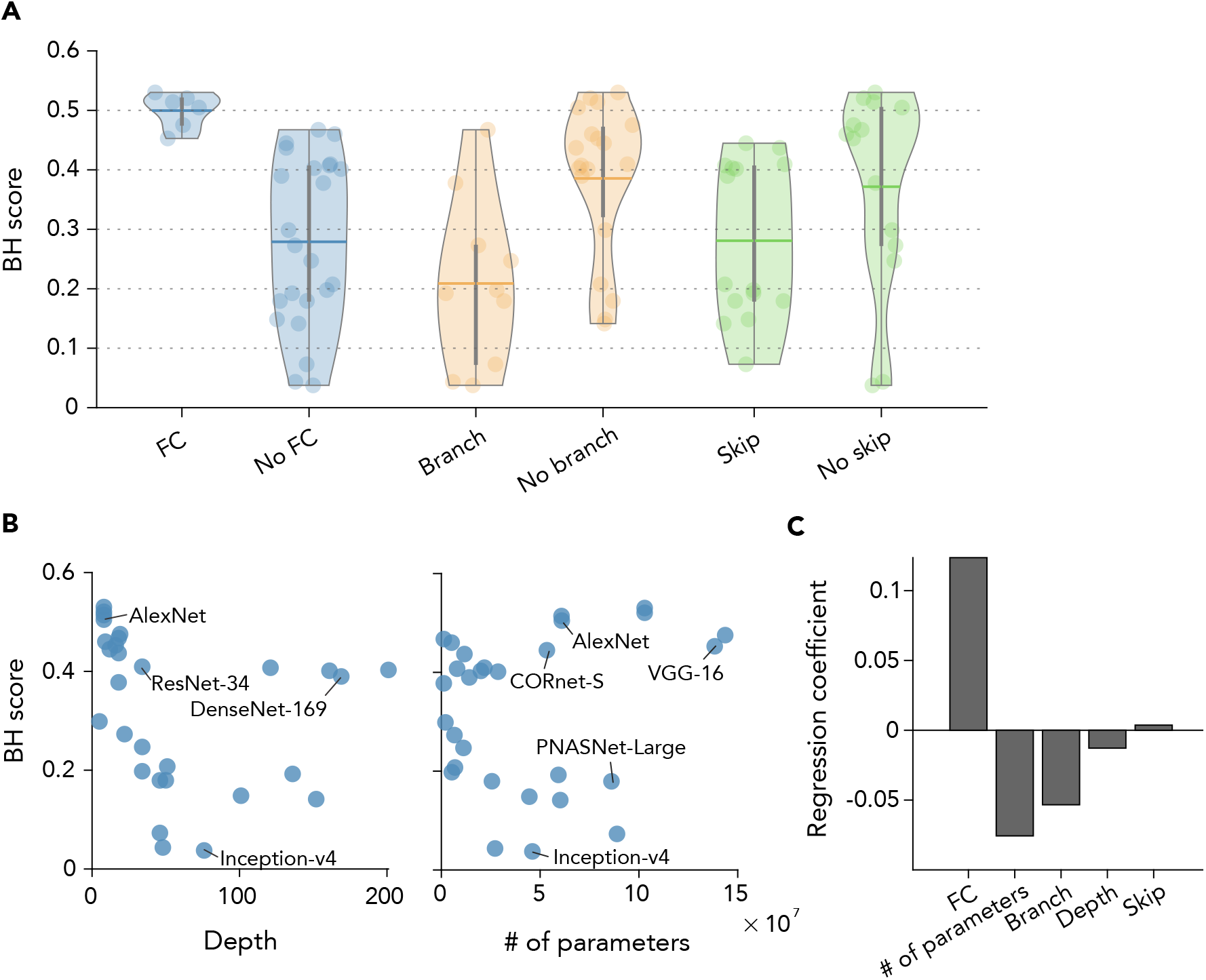
Relationship between BH scores and DNN architectures. **(A)** Comparison between DNNs with and without fully-connected (FC) layers, skip-connections and branch-connections. Thick horizontal lines show the mean BH score across the DNNs. Each dot denotes an individual DNN. Vertical gray lines indicate the interquartile ranges. Shaded areas are the full ranges of data points. **(B)** Relationship to DNN depth and the number of weight parameters. Each Dot denotes individual DNNs. **(C)** Regression analysis. A linear regression model was fitted to explain the BH score with five architectural components. The resultant standardized regression coefficients are shown for each component.

How do FC layers contribute to the high degree of hierarchical homology? As an example, in AlexNet, the distributions of top ROIs for FC layers have peaks at HVC (Figure 7A), leading to a large shift in distribution. To examine whether this tendency is consistently observed for other DNNs with FC layers, we computed the mean top ROI number of each layer and plotted it as a function of the layer number for the 29 DNNs (Figure 7B). Whereas the mean top ROI numbers in the higher layers shifted up to higher visual areas in the DNNs with FC layers, the shift stopped at mid-level visual areas in DNNs without FC layers.

**Figure 7.**
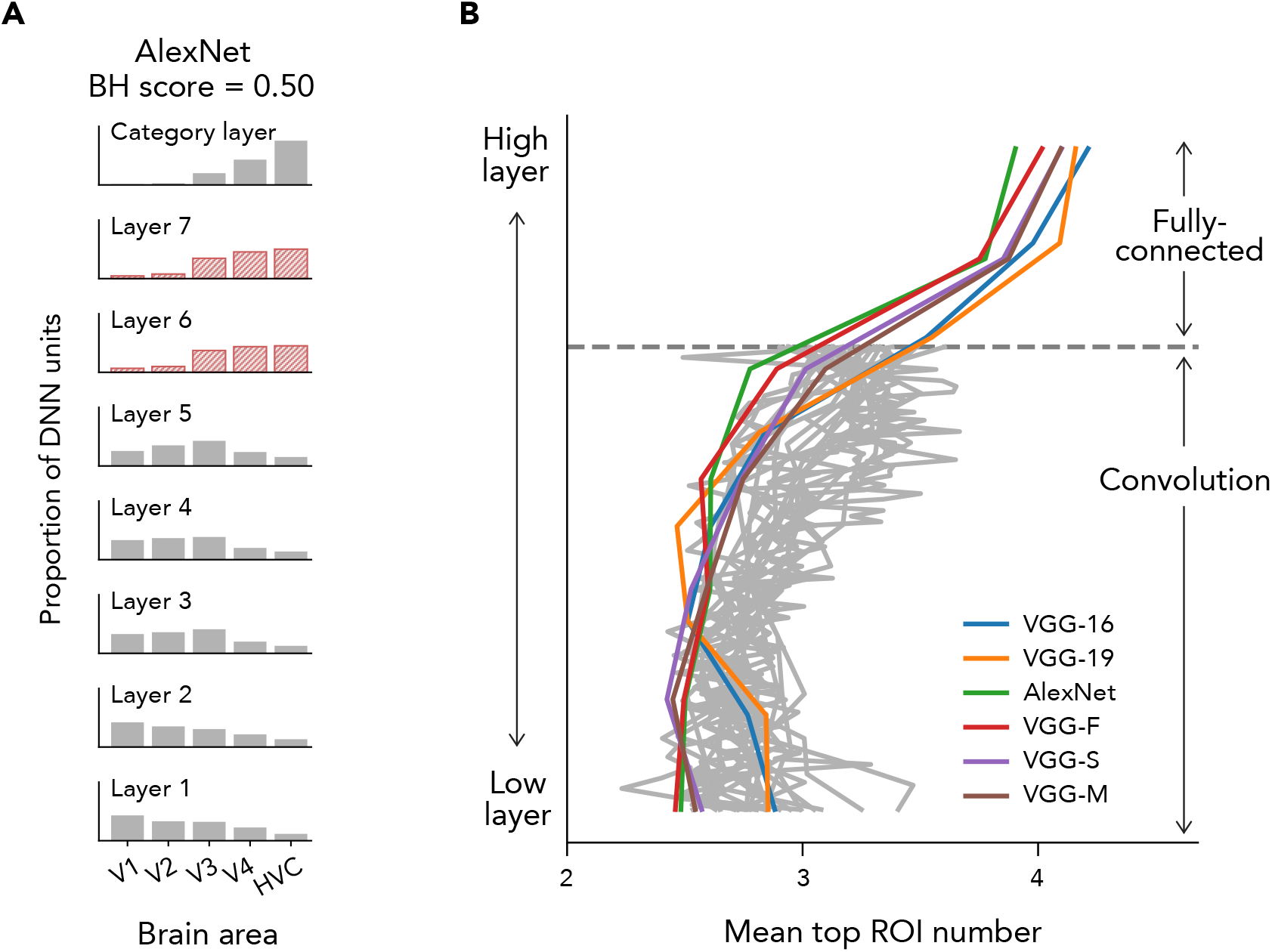
FC layers and the shift of top ROIs. **(A)** The distributions of top ROIs for AlexNet. The distributions for FC layers are shown in red. **(B)** Shifts of the mean top ROIs numbers across all layers for the 29 DNNs. For each DNN, we computed the mean of the distribution of each layer except for the last layer and plotted it as a function of the depth of the DNN layer. The depth of the DNN layer has been rescaled for visualization purposes.

### Experimental manipulation of DNN architecture

To complement the analysis described above, we prepared a base DNN (AlexNet, consisting of five convolutional layers, two fully-connected layers, and one category layer without skip- or branch-connections), and manipulated the number of FC layers from zero to five while maintaining the other architectural characteristics (Transparent Methods: “Manipulation of DNN architecture”). These DNNs were trained on the same ImageNet dataset. The DNN with no FC layers showed a gradual shift of the top ROI distribution up to the second last layer, but had a relatively large gap between the second last layer and the last layer (Figure 8A). In contrast, in DNNs with one, two, and three FC layers, the top ROI distribution gradually shifted over the layers. With even more FC layers, the shift of top ROIs over the convolutional layers became less gradual and the last convolutional layer and the first FC layer exhibited a large gap. Thus, the BH score was highest at the DNNs with two FC layers (Figure 8B). In our comparison of the 29 pre-trained DNNs, all of the DNNs with FC layers (i.e., AlexNet and the VGG family) had two FC layers. This moderate number of FC layers was likely to have produced high BH scores. Convolutional layers and FC layers differ in two main aspects: kernel size and the number of channels. While each unit in an FC layer has connections from all units in the previous layer, units in convolutional layers have spatially limited connections. The spatial range of connections allowed in a convolutional layer is specified by the kernel size. In addition, the number of channels is generally different between convolutional layers and FC layers. The number of channels in an FC layer is typically set to several thousand, whereas that in convolutional layers is set to several hundred. To examine which component is critical for explaining high BH scores of DNNs with FC layers, we tested how each component in FC layers affected the BH score.

**Figure 8.**
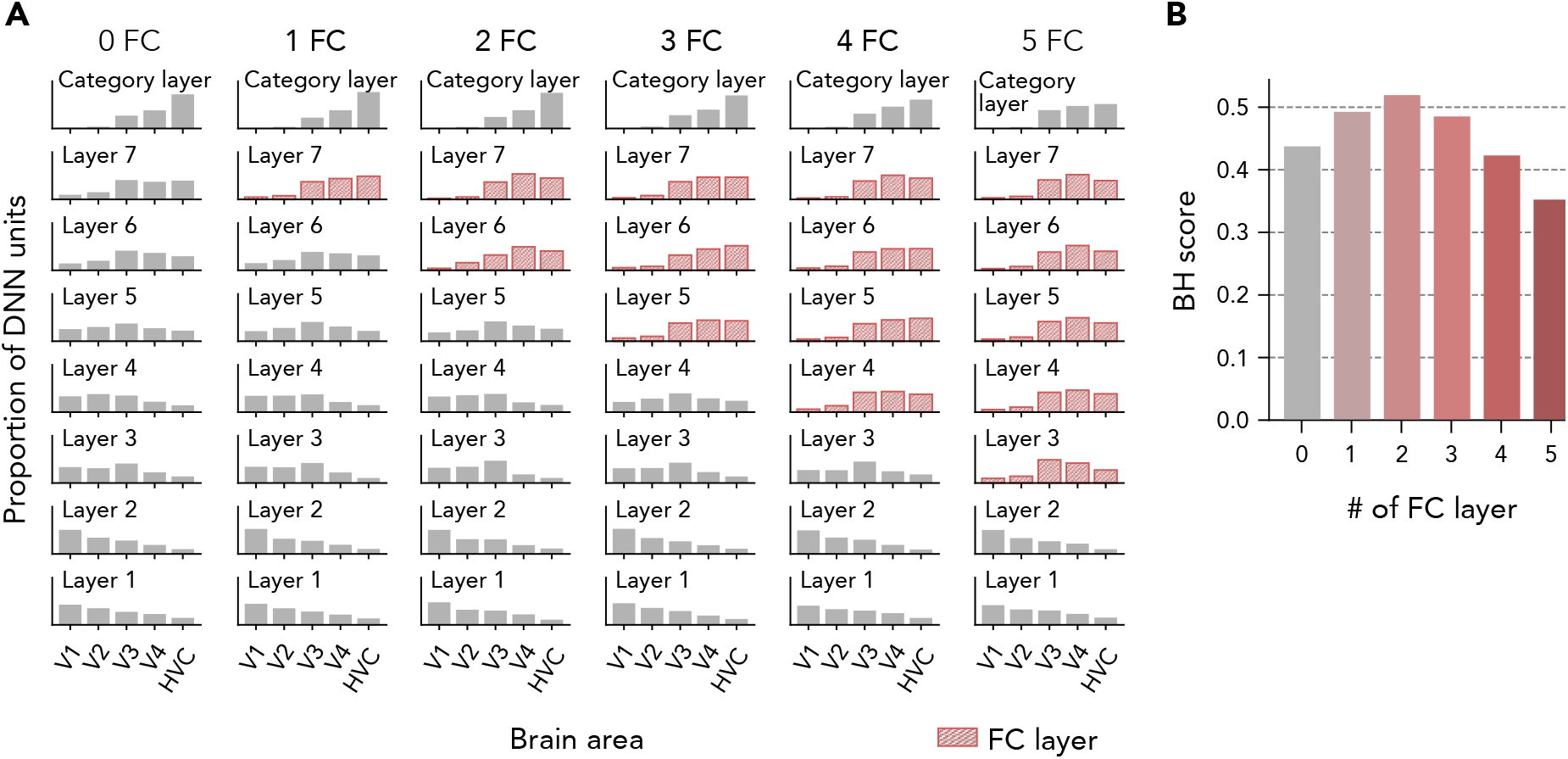
Effect of the number of FC layers. **(A)** Eight-layer DNNs with different numbers of FC layers were trained on ImageNet (see Table 2 for the details of the architectures). The top ROI distributions of top ROIs are shown. FC layers are indicated in red. **(B)** BH scores for DNNs with different numbers of FC layers.

To examine the effects of kernel size, we constructed a DNN with six convolutional layers and one category (fully-connected) layer, and manipulated the kernel size of the last convolutional layer from one to six (Figure 9A; see Transparent Methods: “Manipulation of DNN architecture”). When the kernel size of the DNN is six, each unit in the last convolutional layer has connections from all units in the second last layer. In other words, the last convolutional layer of this DNN is equivalent to a FC layer. By changing this kernel size systematically, we examined how the BH score and the representation of this layer changed depending on this parameter. As the kernel size became larger, the BH score increased (Figure 9B) and the distribution of top ROIs for the last convolutional layer was centered at higher visual areas with some fluctuations (Figure 9C). The results indicate that broad connections in FC layers are a critical factor for brain–DNN homology.

**Figure 9.**
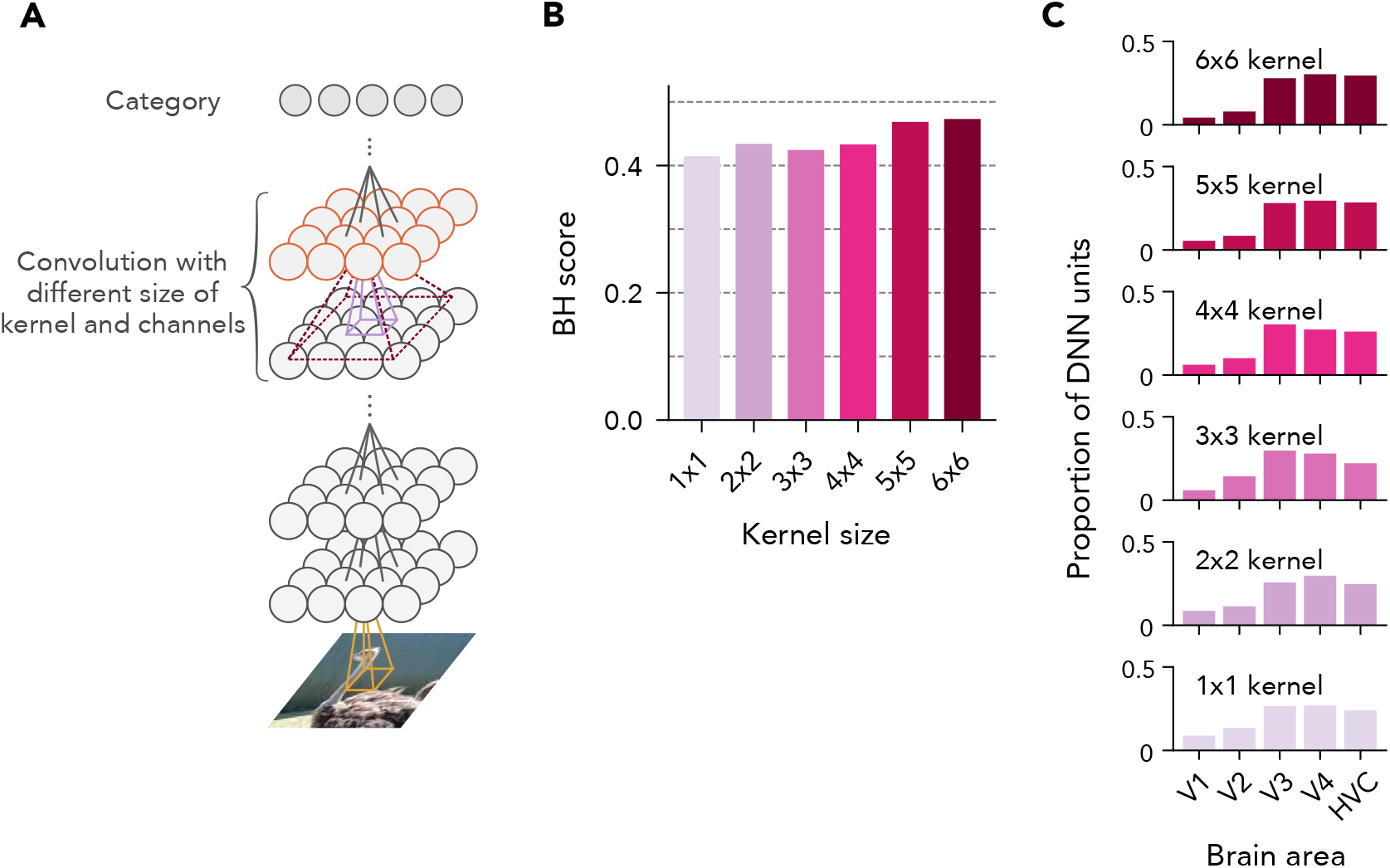
Effects of the width of spatial integration, and the number of channels. **(A)** DNN architecture used for the experiment. We used a seven-layer DNN while changing the kernel size at the second last layer indicated in red (see Transparent Methods: “Deep neural network”). **(B)** BH scores with varied kernel sizes. **(C)** Top ROI distributions at layer 6 with varied kernel sizes.

We also tested how other architectural characteristics (i.e., presence of skip-connections, presence of branch-connections, and depth) affect the BH score by manipulating either of them in base DNNs (AlexNet for the presence of skip- or branch-connections and VGG-16 for depth) (Transparent Methods: “Manipulation of DNN architecture”). Skip- and branch-connections were introduced by replacing convolutional layers with residual blocks of ResNet-18 or inception modules of Inception-v1, respectively. The depth was manipulated by inserting additional convolutional layers. It should be noted that the additional convolutional layers were not included in the calculation of BH scores. Because the number of weight parameters cannot be changed independently, it was not considered here. Consistent with the tendency among the 29 pre-trained DNNs, the distributions of top ROIs for DNNs with skip-connections and branch-connections were relatively flat compared with the base DNN. In addition, the top ROI distributions for the DNN with skip-connections did not monotonically shift from lower to higher visual areas: the centroid of the distribution shifted to higher visual areas at layer 2, then back to lower areas at layer 3 (Figure 10A). As a result, those DNNs showed slightly lower BH scores than the base DNN without them (Figure 10B). When the depth was manipulated, BH scores of deeper DNNs tended to be slightly higher, with sharper peaks. However, the difference was small (Figure 10C, D).

**Figure 10.**
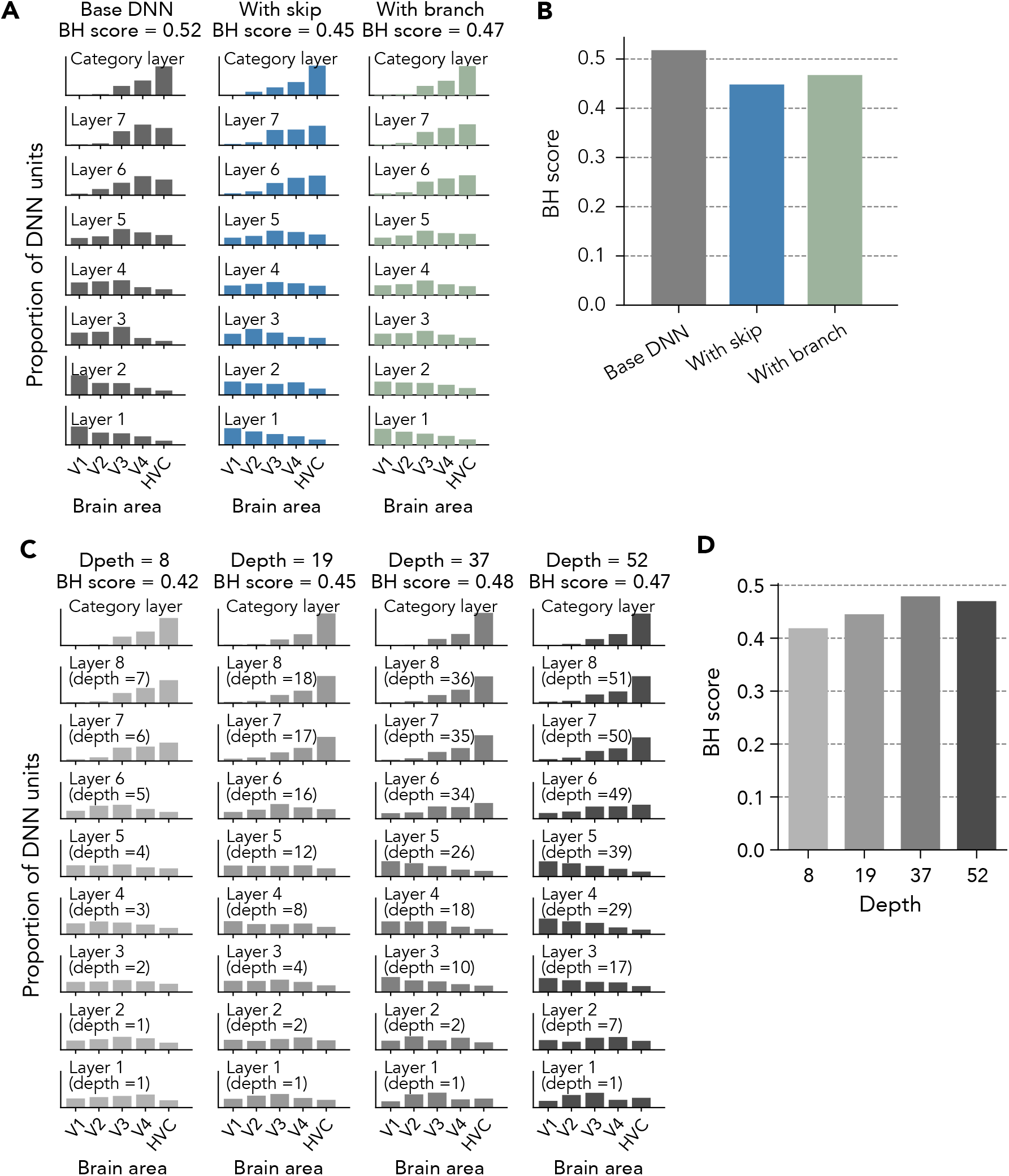
Effects of depth, skip-connections, and branch-connections. **(A)** Effect of skipconnections and branch-connections. A DNN with pure convolutional layers, a DNN with skipconnections and a DNN with branch-connections were prepared and trained on ImageNet using the same procedure (see Transparent Methods: “Deep neural network training” for details of the architectures). **(B)** BH scores for the base DNN, DNN with skip-connections and DNN with branchconnections. **(C)** Effect of depth. Four DNNs with different depths were prepared by inserting convolution layers into each layer. The layer number and depth at each layer are also shown. **(D)** BH scores for DNNs with different depths.

## Discussion

In the current study, we presented a method for quantifying the hierarchical similarity between the human brain and deep neural networks (DNNs) and its applications in an attempt to elucidate the characteristics that make DNNs hierarchically brain-like. We characterized individual DNN units by their best decodable visual areas (top ROIs) in fMRI decoding analysis and quantified the correspondence of hierarchical representations between the brain and DNNs. The distributions of top ROIs revealed differences in the hierarchical development of representations between DNNs (Figure 2). Our proposed metric, the brain hierarchy (BH) score, was negatively correlated with image recognition performance across DNNs (Figure 3), suggesting that high-performance DNNs are not necessarily brain-like. This negative correlation with image recognition performance was consistently observed with different image datasets (Figure S2). Untrained DNNs showed poor BH scores (Figure 4), indicating the importance of DNN training for hierarchical homology with the human brain. The BH score was robust to optional choices of procedures (Figure S3). Complementary encoding analysis also provided similar comparison results across DNNs (Figure S5 and S6). By comparing DNNs with different architectures, we identified architectural characteristics that were associated with the degree of hierarchical homology (Figure 6). DNNs with fully-connected (FC) layers exhibited higher BH scores, and DNNs with skip- and branch-connections exhibited lower BH scores by smaller margins. DNNs with FC layers gradually developed internal representations from those similar to lower visual areas to those similar to higher visual areas, whereas DNNs without FC layers lacked layers whose representations were similar to higher visual areas (Figure 7). This observation was also confirmed by a manipulation experiment (Figure 8). Additional experiments provided further support for the importance of broad spatial integration at FC layers (Figure 9). The presence of skip-connections and branch-connections tended to degrade the degree of hierarchical homology by flattening and non-monotonically swinging the top ROI distributions, respectively (Figure 10).

We have provided code and data to compute the BH score for any DNN (see Transparent Methods: “Data and Software Availability”). Given a DNN, users first obtain the unit activations for the stimulus images used in our functional magnetic resonance imaging (fMRI) experiments. The stimulus images are available on request form (https://forms.gle/ujvA34948Xg49jdn9). Our code repository (https://github.com/KamitaniLab/BHscore) includes scripts for calculating unit activation for the 29 DNNs tested in this study. Users can run the decoding analysis with the DNN unit activations and the fMRI data of individual brain areas (ROIs) shared in a ready-to-analysis format at figshare (https://figshare.com/articles/Deep_Image_Reconstruction/7033577). The scripts for the decoding analysis are available in our code repository. Finally, the BH score is calculated from decoding accuracies by a function in our code repository. Complementary encoding-based procedures can be performed in a similar way.

The BH score is based on the decoding of individual DNN units. Compared with encoding and representational similarity analysis, this method allows us to characterize a DNN at the finest level (Horikawa et al., 2019). In contrast, previous studies based on encoding analysis examined the similarities between a DNN and the brain by predicting brain activations (e.g., neuronal and fMRI responses) from the DNN’s unit activation patterns across thousands of DNN units (Cadieu et al., 2014; Guclu and van Gerven, 2015; Schrimpf et al., 2018; Wen et al., 2018; Yamins et al., 2014; Yamins and DiCarlo, 2016; Khaligh-Razavi and Kriegeskorte, 2014). Likewise, representational similarity analysis has been used to detect the similarities to the brain based on the activation patterns across those units (Jozwik et al., 2019; Khaligh-Razavi and Kriegeskorte, 2014; Storrs et al., 2020). In addition, most of these studies applied dimension reduction techniques to the DNN activation patterns due to their high dimensionality, potentially overlooking fine representations encoded in individual units. Despite these differences, our complimentary assessment based on encoding analysis provided similar comparison results to assessment with the original BH score (Figure S5), suggesting the robustness of the results to the choice of analysis methods.

Our comparative results using the BH score were robust to several choices of procedures. The original BH score was strongly correlated with those computed without the unit exclusion procedure (Figure S3A) or the nonlinear transformation (Figure S3B) across the 29 DNNs. Meanwhile, the raw values of the BH score were systematically affected by these procedures. The unit exclusion procedure tended to increase/decrease the BH score for DNNs with high/low scores, respectively. These procedures generally broadened the range of BH scores, suggesting that they may improve the sensitivity of the BH score.

The BH score was also highly consistent across fMRI datasets from different subjects (Figure S4), consistent with our previous study showing high correlations of the decoding accuracies of individual DNN units between different subjects (Horikawa et al., 2019). While the BH score showed high consistency across normal subjects, brain disorders such as schizophrenia have recently been proposed to be associated with disorganized object representations in the brain (Nishida et al., 2020). The BH score based on fMRI responses from participants with such disorders may reveal the functional differences compared with typical brains.

In the current study, we found that the BH score was negatively correlated with image recognition performance across the 29 pre-trained DNNs (Figure 3). Thus, high-performance DNNs do not necessarily exhibit hierarchical representations that are similar to the brain. Although early work suggested that DNNs with improved recognition performance are likely to provide better computational models of the brain (Yamins and DiCarlo, 2016), a recent study (Schrimpf et al., 2018) reported that image recognition performance and brain–DNN similarity was more weakly correlated in recently developed high-performance DNNs (DNNs with ≥ 70% ImageNet top-1 accuracy). Our comparative results using the BH score also showed a dissociation between image recognition performance and similarity to the brain. As one of the gaps in the object recognition process between DNNs and the brain, a recent computer vision study suggested that DNNs trained on ImageNet tended to classify object images according to their textures, whereas humans classify object images based on their shapes (Geirhos et al., 2019). A subsequent study quantitatively compared this texture bias between AlexNet and ResNet-50, and showed that the global-pooling operation in ResNet-50 largely removes shape information and strengthen the texture bias (Hermann and Kornblith, 2019). Although both DNNs have a strong texture bias, AlexNet preserves relatively richer shape information in their FC layers. This suggests that FC layers mitigate DNNs’ texture bias and make processing in the higher layers more brain-like, which may explain our comparative results between DNNs with and without FC layers. In contrast, because FC layers have large numbers of weight parameters and increase the risk of overfitting, global-pooling operations are more commonly adopted in recent high-performance DNNs. Likewise, elucidating the differences between high-performance DNNs and the brain remains an important challenge, which if solved, would provide important insights into the properties of DNNs.

In addition to the quantification of the brain–DNN hierarchal homology with the BH score, we characterized individual DNN layers by the distributions of their top ROIs. The distributions of top ROIs tended to show specific patterns depending on the architectures of DNNs. DNNs with FC layers (e.g., AlexNet and the VGG family) showed unimodal and sharp distributions at the layer. The peak of the distribution was monotonically shifted from lower to higher visual areas along with the hierarchy of the DNN layers, leading to high BH scores for those DNNs. In contrast, DNNs with branch-connections (e.g., the Inception family, NASNet and PNASNet) tended to have flat distributions. Because the branch-connections develop their features through parallel convolutions with different kernel sizes, representations corresponding to different visual areas may be mixed into single concatenated layers, flattening the distributions. In the DNNs with skip-connections (e.g., the ResNet family, the DenseNet family, NASNet, and PNASNet), the peak of the distribution tended to oscillate between V1 and V3, possibly reflecting the bypassing of representations between lower and higher layers. Although brain regions are connected not only by feedforward path but also by branch-like or skip-like connections (Felleman and Van Essen, 1991; Sporns and Zwi, 2004), DNNs with such connections do not yield hierarchical representations similar to the brain. Instead, simple feedforward DNNs acquire more hierarchically similar representations to the brain, suggesting that feedforward connections among brain regions play a dominant role in forming hierarchical representations in the brain.

While deeper DNNs tended to have lower BH scores among the 29 pre-trained DNNs (Figure 6B), the opposite tendency was observed in the DNNs trained after only their depth was manipulated (Figure 10C, D). Many of the deeper pre-trained DNNs had skip-connections in their architectures. Skip-connections are often used to mitigate the risk of gradient vanishing in very deep networks (He et al., 2015). The covariation of these architectural factors in the 29 pretrained DNNs may account for the discrepancy in the effect of depth found in the manipulation experiment.

Our results suggest that full connections in the last few layers (FC layers) make the representations similar to those in the higher visual areas and thus lead to greater hierarchical homology (Figure 8). FC layers can spatially integrate visual features to achieve translation-invariant representation of object categories. fMRI activity in the higher visual areas, including lateral occipital complex (LOC), the fusiform face area (FFA), and the parahippocampal place area (PPA), is associated with processing translation-invariant information of object categories (Carlson et al., 2011). Thus, spatial integration of local visual features may play a common crucial role in hierarchical development of spatially invariant visual representation in both DNNs and the brain. In addition, a recent study reported that DNNs with FC layers exhibit better generalizability across datasets than all convolutional DNNs (i.e., DNNs without FC layers) (Zhang et al., 2018). FC layers may also play a critical role in achieving human-level generalizability. In contrast, all convolutional DNNs tend to show better ImageNet top-1 accuracies than DNNs with FC layers. This is presumably because all convolutional DNNs have smaller numbers of weight parameters due to weight sharing, allowing for efficient learning given a limited amount of training data. Thus, only performance-optimization for a specific task may not lead to brain-like DNNs.

We found that DNNs with random weights (i.e., untrained DNNs) showed markedly lower BH scores than DNNs trained for the classification task on ImageNet (Figure 4). Similarly, DNNs trained for object classification have been shown to have a more similar representation to human IT than untrained DNNs (Storrs et al., 2020). These results suggest that goal-driven task optimization of DNNs is critical for brain-like representations. However, another recent study reported that DNNs trained in an unsupervised manner could explain neuronal responses in monkey V1, V4, and IT cortex to a comparable degree to DNNs trained for object classification (Zhuang et al., 2020). It remains unclear whether task-optimization (i.e., supervised learning) is necessary, or if just “experience” is sufficient for DNNs to acquire the hierarchical homology.

Although we quantitatively characterized the hierarchical homology of visual representations between DNNs and the human brain, the BH score could be used for examining hierarchical homology of representations in different modalities (e.g., auditory representations, tactile representations) or that between DNNs and the brain of other species. Previous studies examined the hierarchical homology between a DNN trained for sound classification and the human auditory cortex (Kell et al., 2018) and the hierarchical homology between VGG16 and the mouse visual cortex (Cadena et al., 2019). Another study proposed several types of biologically-feasible DNN that imitate hierarchical representations of the rodent tactile system (Zhuang et al., 2017).

It remains unclear which type of DNN best captures the properties of real rodents. The BH score would provide a quantitative tool for comparison of different types of DNN in terms of the similarity to given hierarchical representations in those modalities and species.

## Limitations of the study

The BH score is only applicable to single-path sequential hierarchies of deep neural networks (DNNs) and brains. For the formalization of hierarchy, we assumed that the hierarchy is represented by an ordinal scale (i.e., layer and ROI numbers). Thus, the BH score does not incorporate multi-path or non-sequential hierarchical structures such as branching, collateral path, or loop (recurrence). Although some DNNs tested in this study have branch-, skip-, or recurrent-connections, we selected the output layers of the branch-, skip-, or recurrent-blocks (submodules) instead of regarding individual layers in the block as multiple layers. Thus, the hierarchy of DNNs was summarized as a single feedforward pathway. In the current study, we focused on DNNs developed for image recognition and the brain regions underlying object recognition (i.e., the ventral visual pathway), in which neural representations are assumed to develop through a single pathway. To quantify the homology between more complex DNNs (e.g., multi-path, and/or recurrent neural networks) and the whole brain networks, a more sophisticated index will be required.

The choice of brain regions (ROIs) and their hierarchical order are critical for the BH score; different sets and orders of ROIs produce different scores in the same DNN. In this study, we selected brain regions in the ventral visual pathway (V1, V2, V3, V4, and higher visual cortex [HVC]), which underlies object recognition. The vital role in object recognition and the structural and functional hierarchy of these areas have been well established in neuroscience research, supporting the notion that that our set and ordering of ROIs captured the hierarchy of object recognition. Nevertheless, this ROI selection method based on known functional anatomy is inevitably user-dependent and has the potential to scatter BH scores. Moreover, the prior knowledge-based selection of ROIs has a potential inherent risk of overlooking hierarchy in the brain that is not incorporated in our prior knowledge. Instead of such prior knowledge-based ROIs, brain hierarchy characterized by data-driven approaches (e.g., Margulies et al., 2016) can be used for the assessment of BH scores and could yield alternative measures.

As architectural characteristics of interest, we did not focus on the presence of recurrent connections because only one DNN has recurrent connections among the 29 DNNs examined in the current study (Table 1). We limited the DNNs tested here to those trained on the same ImageNet classification task for fair comparison, and there were few available pre-trained DNNs with recurrent connections satisfying this limitation. However, several studies recently developed DNNs with recurrent-connections, suggesting that those DNNs capture the dynamics of neuronal responses in the ventral visual areas (Nayebi et al., 2018; Spoerer et al., 2017). Testing new DNNs designed to imitate the processing in the cortex will be helpful for elucidating the importance of recurrent connections in the human visual system.

## Acknowledgments

The authors would like to thank Tomoyasu Horikawa, Guohua Shen, Fan Cheng, and Mohamed Abdelhack for helpful comments on the manuscript. The data used in the study were collected using the MRI scanner and related facilities of Kokoro Research Center, Kyoto University. This work was supported by The New Energy and Industrial Technology Development Organization (NEDO), JSPS KAKENHI Grant Numbers JP15H05920, JP15H05710, and AMED under Grant Number JP18dm0107151.

## Author contributions

Conceptualization, Y.K.; Methodology, S.N. and K.M.; Validation, S.N., K.M., and S.C.A.; Formal Analysis, S.N., K.M., and S.C.A.; Investigation, S.N., K.M., and S.C.A.; Resources, Y.K.; Writing – Original Draft, S.N., K.M., and S.C.A.; Writing – Review & Editing, S.N., K.M., S.C.A., and Y.K.; Visualization, S.N., K.M., and S.C.A.; Supervision, Y.K.; Funding Acquisition, Y.K.

---

**Figure S1.**
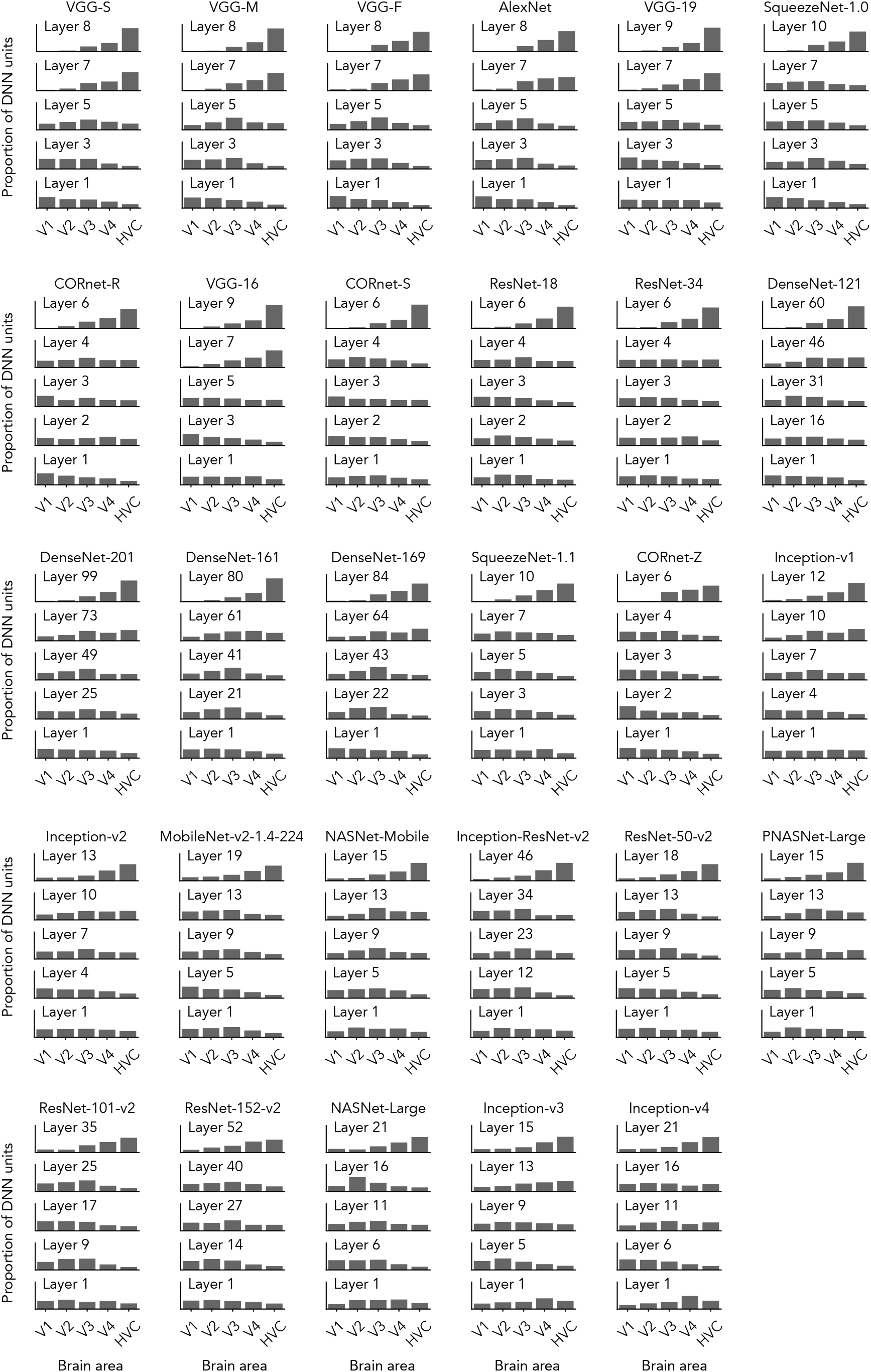
Top ROI distributions for all 29 deep neural networks (DNNs), **Related to Figure 2.** The distributions of top ROIs for individual layers of 29 DNNs are shown. For visualization purposes, the results for five layers sampled from each DNN are plotted.

**Figure S2.**
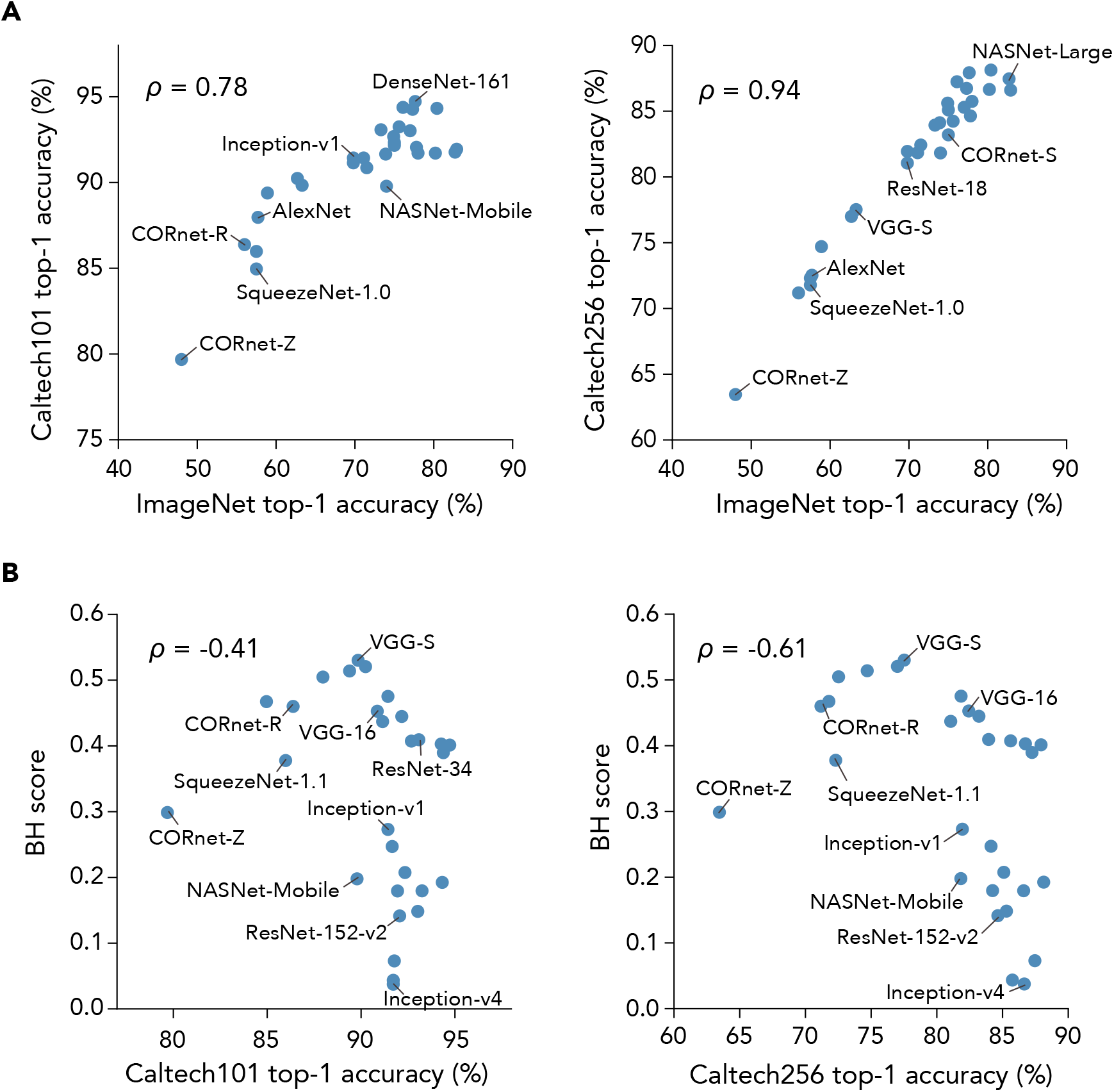
Image recognition performance of DNNs evaluated using different image datasets, **Related to Figure 3. (A)** Image recognition performance. ImageNet top-1 accuracy is plotted against the image recognition accuracy evaluated on Caltech-101 (left) and Caltech-256 (right). To evaluate the accuracy on Caltech-101 and Caltech-256, a multinomial logistic regression classifier was trained using the second last layer of each DNN as input. **(B)** The brain hierarchy (BH) scores and image recognition accuracies on Caltech-101 and Caltech-256. The BH scores for 29 DNNs are plotted against the image recognition accuracies evaluated on Caltech-101 and Caltech-256.

**Figure S3.**
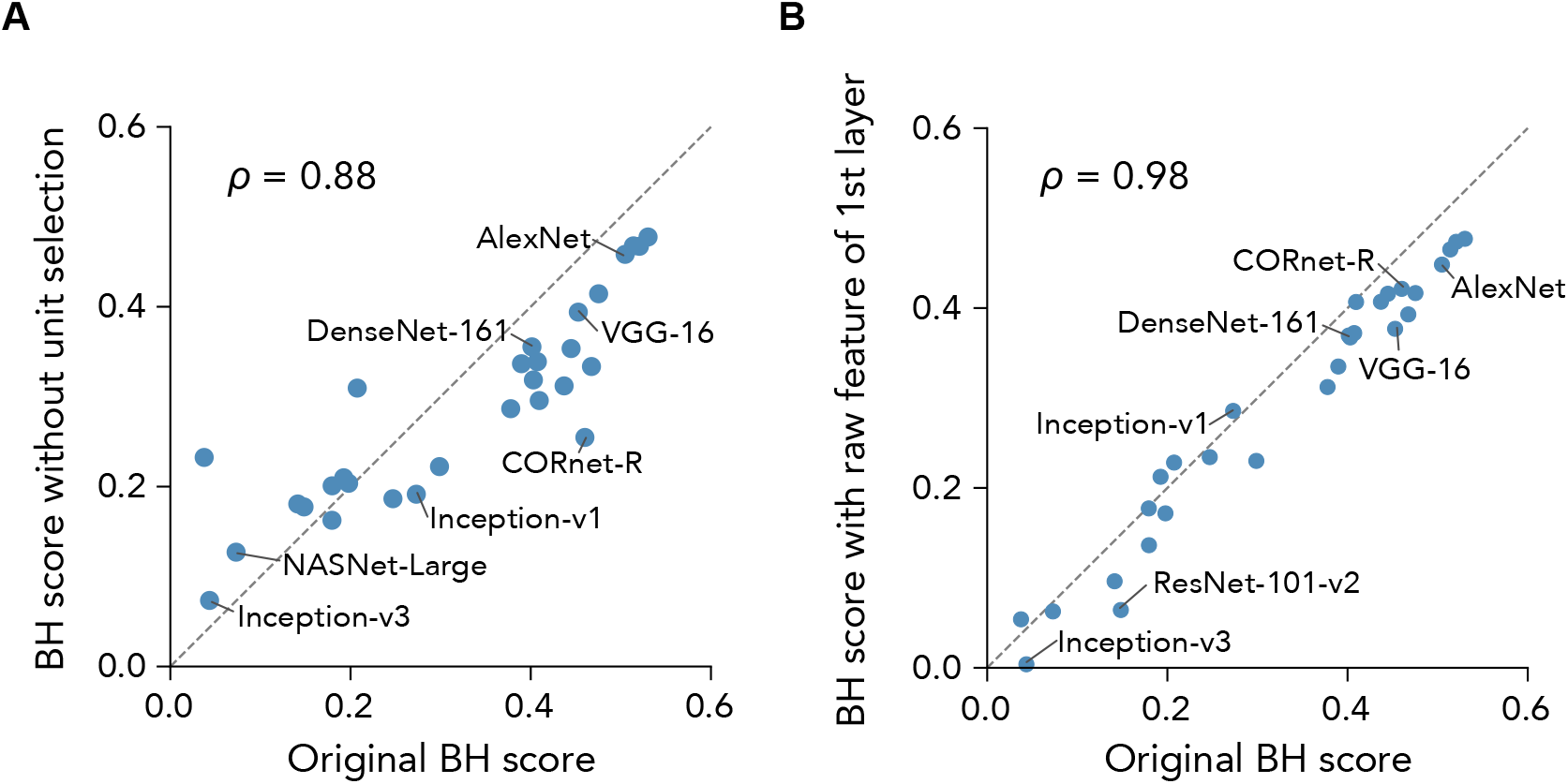
Robustness of BH scores on computation procedures, **Related to Figure 3. (A)** BH scores computed with and without unit selection. BH scores computed without unit selection are plotted against BH scores computed with our original definition of the BH score. **(B)** BH scores computed with raw features of the earliest layer and with our original definition of the BH score. BH scores computed with raw features of the earliest layer are plotted against BH scores computed with our original definition of the BH score.

**Figure S4.**
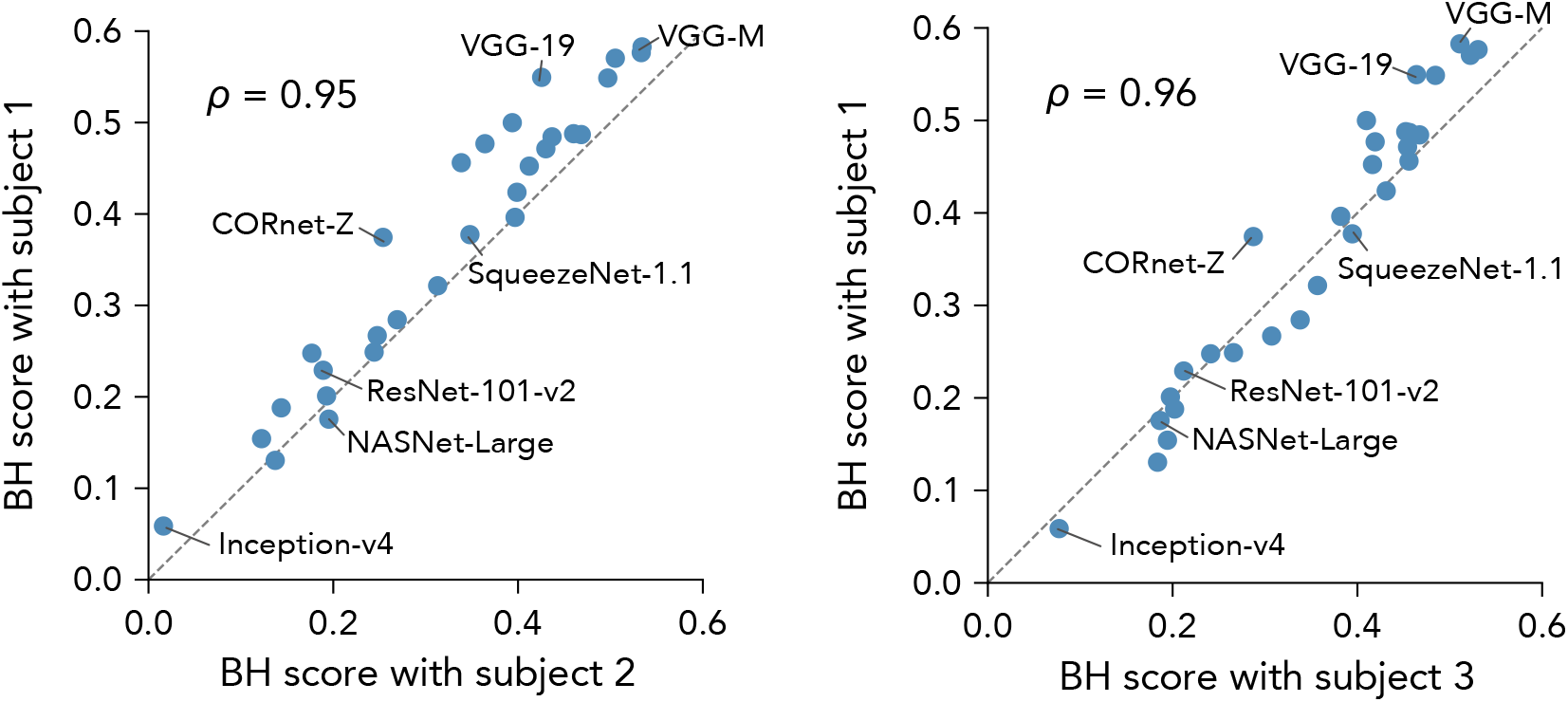
Consistency of the BH scores across fMRI data from different subjects, **Related to Figure 3.** To confirm the robustness of BH scores, scores were computed from fMRI data of each single subject. BH scores computed from the functional magnetic resonance imaging (fMRI) data of one subject are plotted against BH scores computed from the fMRI data of other subjects.

**Figure S5.**
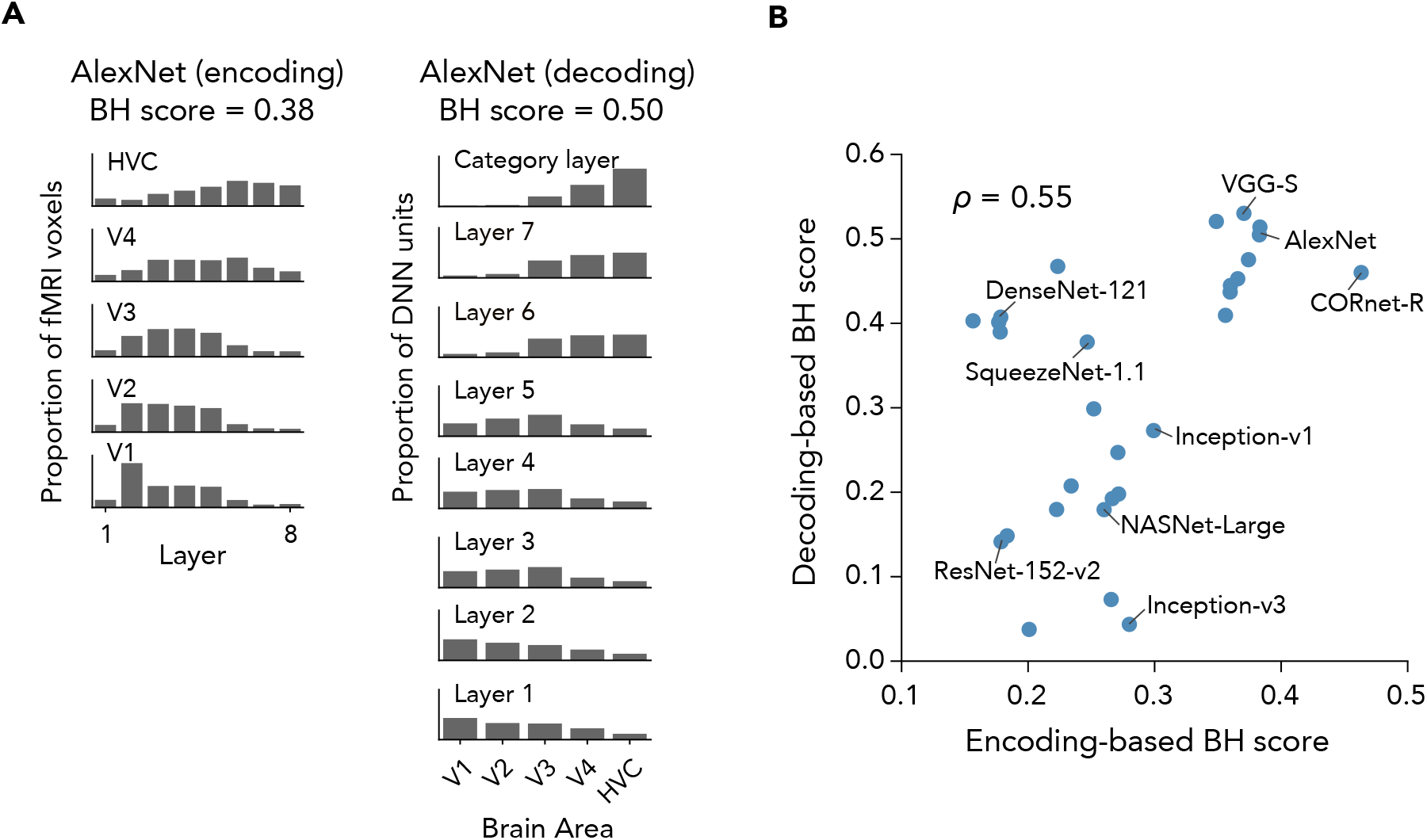
Consistency between BH scores obtained by decoding analysis and BH scores obtained by encoding analysis, **Related to Figure 3. (A)** Distributions of best predicting DNN layers obtained by encoding analysis and decoding analysis. Encoding analysis was performed where the fMRI responses of individual voxels were predicted from the responses of each DNN layer. For each fMRI voxel, the DNN layer showing the highest prediction accuracy (best layer) was assigned. The distributions of best layers for individual ROIs (left) and distributions of top ROIs for individual layers of AlexNet (right) are shown. **(B)** BH scores obtained by decoding analysis and by encoding analysis. BH scores obtained by decoding analysis and by encoding analysis are shown in a scatter plot.

**Figure S6.**
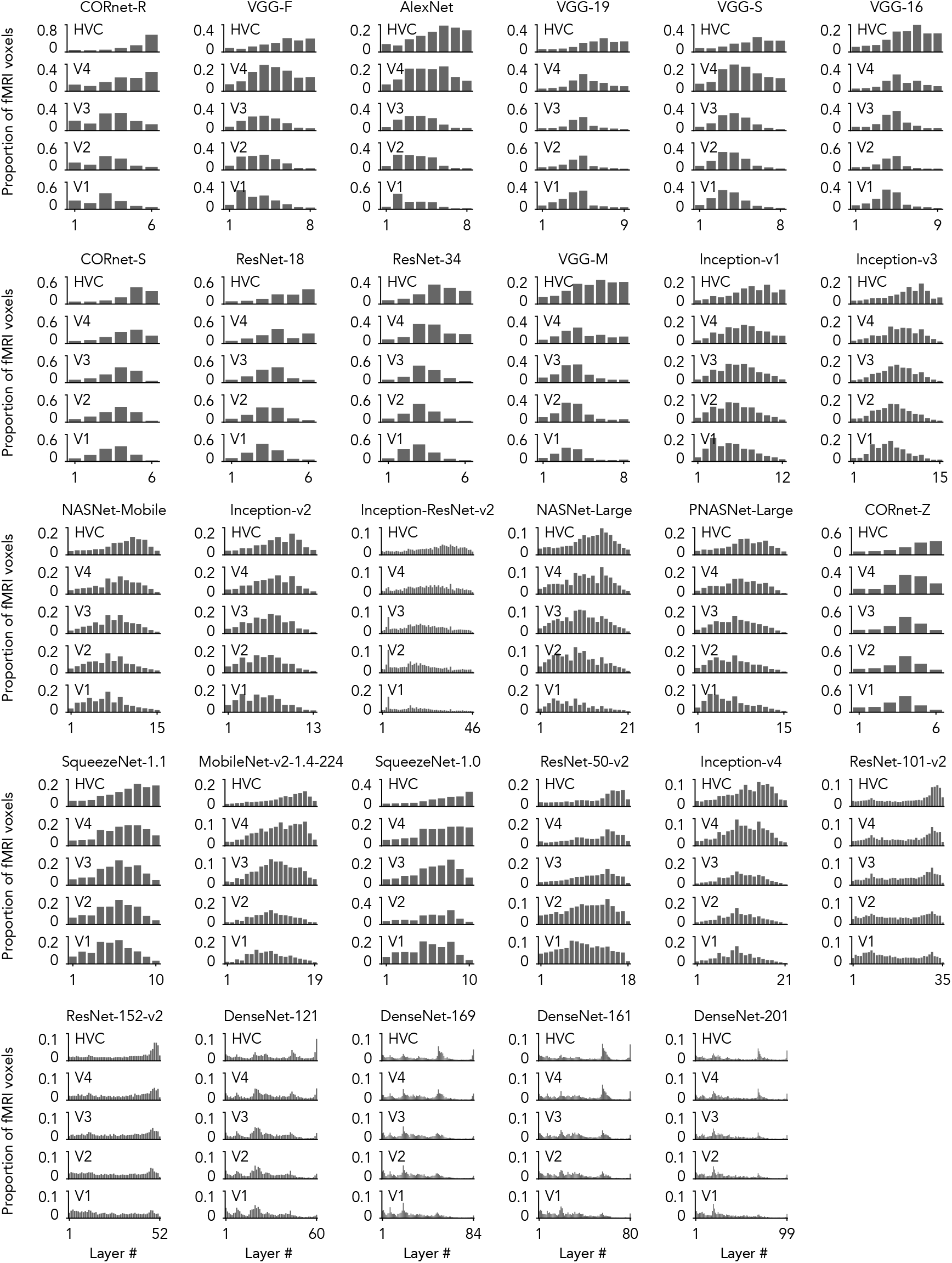
Distributions of top DNN layers for all 29 DNNs, **Related to Figure 3.** In the encoding analysis, fMRI responses of individual voxels in each ROI were predicted from the DNN unit responses in each DNN layer of 29 DNNs. Then, for each voxel, the DNN layer showing the highest prediction accuracy among the layers in each DNN (top layer) was identified. The distribution of the best layers for voxels in each ROI is shown. The range of the y-axis is changed for visualization purposes.

**Figure S7.**
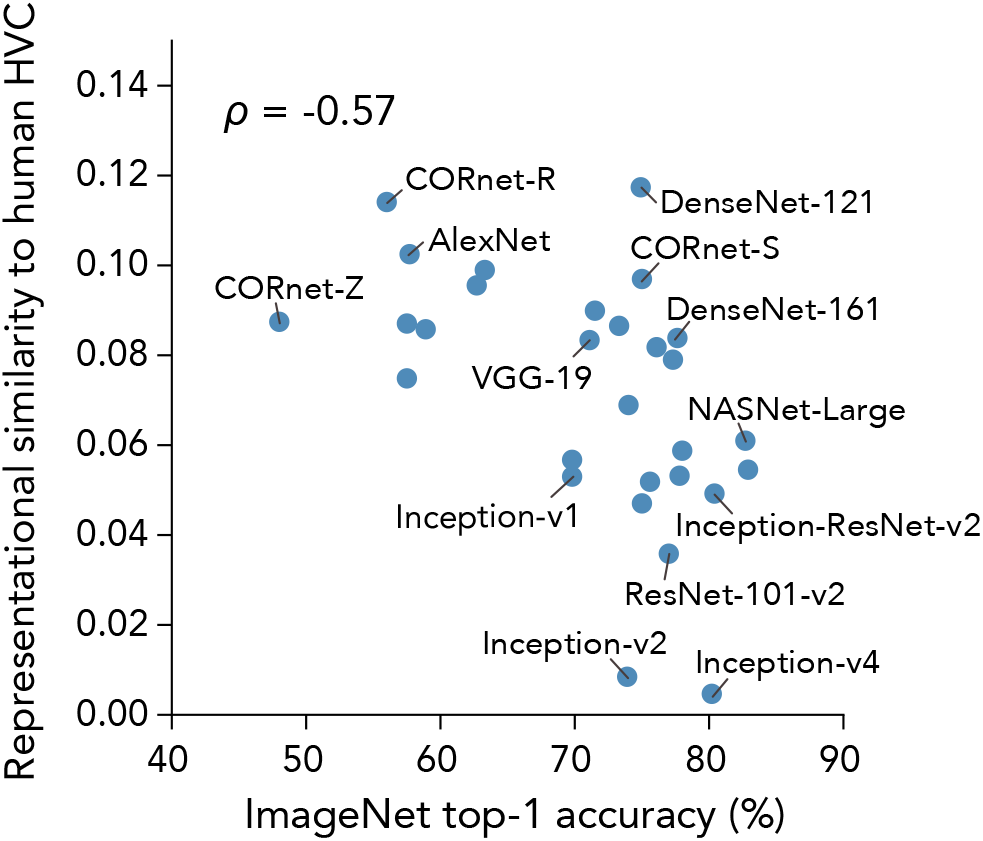
Representational similarity to human higher visual cortex (HVC) and ImageNet top-1 accuracy, **Related to Figure 3.** The representational similarities between DNNs and the human HVC were quantified by following the procedure reported by Jozwik et al. (2019), and those are plotted against ImageNet top-1 accuracies

## Transparent Methods

### fMRI dataset

We used the functional magnetic resonance imaging (fMRI) dataset collected in Shen et al. (2019). The dataset, which is publicly available at OpenNeuro (https://openneuro.org/datasets/ds001506), includes fMRI data from three human subjects. During fMRI scanning sessions, subjects viewed natural object images selected from ImageNet (2011, fall release) (Deng et al., 2009). The fMRI experiment was composed of two types of sessions: training and test sessions. In training sessions, 1200 images selected from 150 categories were used. Each image was presented five times. In test sessions, 50 images selected from 50 categories were presented. Each image was presented 24 times. During the experiment, subjects performed one-back repetition tasks. Voxel size was 2×2×2 mm and TR was 2 s in the fMRI measurement. The fMRI responses in the training and test sessions were used for training and test of decoders/encoders, respectively (see **Decoding analysis** and **Encoding analysis**).

Motion correction and anatomical-functional coregistration to individual brains were performed on the fMRI signals with SPM (http://www.fil.ion.ucl.ac.uk/spm). After preprocessing, nuisance parameters (head motion parameters and linear trend) were regressed out from the signal of each voxel. Then, the signal amplitudes were normalized relative to the mean amplitude during the initial rest period (24 s) of each run, and despiked by reducing extreme values (beyond ±3 standard deviations in each run). The signal time series were shifted by 4 s to compensate for hemodynamic delays. The fMRI response to each image was obtained by averaging fMRI signals during the presentation block (8 s) of each image.

### Region of interest

Five visual areas (V1, V2, V3, V4, and higher visual cortex [HVC]) were included in the analysis. All regions of interest (ROIs) were defined functionally on individual brains. V1, V2, V3, and V4 were delineated based on standard retinotopy experiments (Engel et al., 1994; Sereno et al., 1995). The HVC was manually delineated as a continuous region covering the lateral occipital complex (LOC), the fusiform face area (FFA), and the parahippocampal place area (PPA). The LOC, FFA, and PPA were identified by the conventional functional localizer experiments (Epstein and Kanwisher, 1998; Kanwisher et al., 1997; Kourtzi and Kanwisher, 2000).

### Deep neural networks (DNNs)

We compared 29 pre-trained DNNs in this study (Table 1). All DNNs were pre-trained on ImageNet ILSVRC 2012 dataset (Deng et al., 2009) to classify given images into 1000 object categories. We used AlexNet (Krizhevsky et al., 2012), VGG-16, VGG-19 (Simonyan and Zisserman, 2014)^a^, VGG-S, VGG-M, VGG-F (Chatfield et al., 2014)^b^, ResNet-18, ResNet-34 (He et al., 2015)^c^, ResNet-50-v2, ResNet-101-v2, ResNet-152-v2 (He et al., 2015)^d^, Inception-v1, Inception-v2, Inception-v3, Inception-v4 (Deco et al., 2015; Szegedy et al., 2016, 2014)^e^, Inception-ResNet-v2 (Szegedy et al., 2016)^f^, SqueezeNet-1.0, SqueezeNet-1.1 (Iandola et al., 2016)^g^, DenseNet-121, DenseNet-161, DenseNet-169, DenseNet-201 (Huang et al., 2016)^h^, NASNet-Mobile, NASNet-Large (Zoph et al., 2018), PNASNet-Large (Liu et al., 2018)^i^, MobileNet-v2-1.4-224 (Sandler et al., 2019)^j^, CORnet-Z, CORnet-R, and CORnet-S (Kubilius et al., 2018)^k^.

To characterize hierarchy in DNNs, we included several representative layers of each DNN in the analysis: the first layer, the last fully-connected (FC) layer (referred to as the “category layer” in this study), the other FC layers, all convolutional layers in DNNs without submodules (i.e., AlexNet, VGG-S, VGG-M, and VGG-F), and the output layers of submodules (i.e., convolutional, residual, or inception blocks) in DNNs with submodules. For residual blocks, we regarded the sum of skip layers as the output of the block. Hereafter, “layer” means the representative layers unless otherwise stated.

### Decoding analysis

For each individual unit in a DNN, we constructed a decoder to predict (decode) the unit activation to an image from fMRI response patterns to the same image. We used linear regression with L2-regularization to construct the decoders. The input feature of the decoder was an fMRI response pattern of 500 voxels in one of the five ROIs. We selected the 500 voxels that showed the highest absolute correlations between their fMRI signals and the target unit activations in the training sessions (Shen et al., 2019). The unit activations in the first layers of DNNs were converted into absolute values, since both increments and decrements of stimulus luminance strongly modulate fMRI signals in the early visual cortex (Haynes et al., 2004) and the absolute values of unit activations in the first layer are better predicted than raw values (Shen et al., 2019). The decoders were trained with fMRI data in the training sessions for each subject. After the training of decoders, we predicted activations of the individual units from fMRI response patterns in the test sessions. In the test fMRI data, responses of individual voxels to the same images were averaged across trials to increase signal-to-noise ratio of the fMRI signals. Thus, we obtained for each DNN unit 50 predicted activation values corresponding to 50 images in the test sessions. The prediction accuracy was evaluated as the Pearson correlation coefficient between the actual and predicted unit activations across the test images. For each DNN layers, activations of randomly selected 1000 units were predicted if the number of units was more than 1000. Otherwise, activations of all unit were predicted. We ran decoding of individual DNN unit activations using fMRI response patterns from five ROIs in three subjects independently.

### Encoding analysis

For individual voxels in the ROIs, we constructed an encoding model to predict the voxel response to an image from unit activation patterns in a DNN layer to the same image. We used linear regression with L2-regularization to construct the encoding models. The input features of the encoding models were activation patterns of DNN units in a layer. For each layer, activations of 500 units were selected in the same procedure as the decoding analysis. As in the decoding analysis, the unit activations in the first layer of DNNs were converted into absolute values. The encoding models were trained with fMRI data in the training sessions for each subject. After the training of encoding models, we predicted responses of the individual voxels from DNN unit activation patterns to the images in the test sessions. Thus, for each voxel, we obtained 50 predicted response values corresponding to 50 images in the test sessions. In the test fMRI data, the responses of individual voxels to the same images were averaged across trials. The prediction accuracy was evaluated as the Pearson correlation coefficient between the predicted and observed responses of voxels across the test images.

In the replication analysis of Schrimpf et al. (2018), the median of encoding accuracies across voxels in V4 and HVC was computed for each DNN layer. The encoding accuracies were obtained for fMRI data of each subject, then averaged across subjects. For a given DNN, the highest encoding accuracy for V4 among the DNN layers was defined as “fMRI V4 encoding accuracy” and the highest mean encoding accuracy for HVC among the DNN layers was defined as “fMRI HVC encoding accuracy”.

### The brain hierarchy (BH) score

For calculation of the brain hierarchy (BH) score, the ROIs were assigned numbers as 1 to 5 from the lower to the higher visual areas (i.e., V1 (1), V2 (2), V3(3), V4 (4), HVC (5)). Similarly, the DNN layers in each DNN were assigned numbers from1 to *N* in order from input to output (*N* is the number of layers in the DNN).

The BH score was calculated for each DNN using the following procedure. We randomly selected three layers from the representative layers except the first and last (category) layers. The five layers (the selected three layers, the first layer, and the last layers) were included in the calculation of the BH score. The layer selection and subsequent calculation of the BH score was repeated 100 times for each DNN. For each unit in the selected layers, we identified the ROI that had the highest prediction accuracy (“top ROI”) based on the decoding analysis. Then, we applied an optional unit selection; units in which prediction accuracy of the top ROI was not significantly higher than the chance level (*t*-test, *p* < 0.05, uncorrected) were excluded from the further analysis. The remaining units were pooled across the DNN layers and subjects. The BH score of a DNN was defined as the Spearman rank correlation coefficient between the layer number and the top ROI number across units in the DNN. To compute the Spearman rank correlation coefficient, the layer numbers and the top ROI numbers were transformed into fractional ranks:

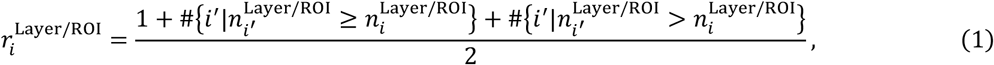

where 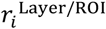 is the fractional rank of the layer number/ROI number of *i*-th DNN unit, 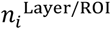 is the layer/ROI number, #{·} denotes the member in the set {·}. The BH score was computed as

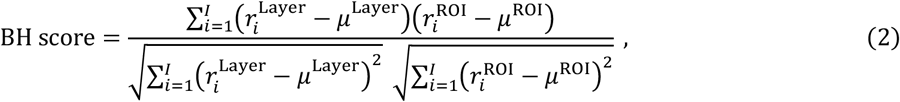

where *μ*^Layer^ and *μ*^ROI^ are the means of 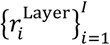 and 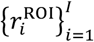, respectively. The encoding-based BH score was computed for each DNN by swapping units and voxels in the calculation of the decoding-based BH score. Unlike the decoding-based BH score, the all representative layers were included in the calculation of the encoding-based BH score. Based on the encoding analysis, we identified for each voxel the layer that had the highest prediction accuracy (“top layer”). We then applied optional voxel selection; voxels in which the prediction accuracy of the top layer was not significantly higher than the chance level (*t*-test, *p* < 0.05, uncorrected) were excluded from the further analysis. The survived voxels were pooled across the ROIs and subjects. The BH score of a DNN was defined as the Spearman rank correlation coefficient between the ROI number and the top layer number across voxels. To compute the Spearman rank correlation coefficient, the ROI numbers and the top layer numbers were transformed into fractional ranks by equation (1). The encoding-based BH score was computed by equation (2).

### Manipulation of DNN architecture

We manipulated the number of FC layers in AlexNet (Krizhevsky et al., 2012) by replacing the layers with convolutional or FC layers. The default AlexNet had five convolutional, two FC, and one category layers. All and the first FC layers were replaced with convolutional layers to create a DNN with zero and one FC layers, respectively. The last one, two, and three convolutional layers were replaced with FC layers to create a DNN with three, four, and five FC layers. DNNs with 0, 1, 2, 3, 4, and 5 FC layers achieved 0.55, 0.57, 0.56,0.51, and 0.46 ImageNet top-1 accuracies, respectively.

To manipulate kernel sizes, we modified a DNN with six convolutional and one category layers. The kernel sizes of the convolutional layers were changed from 1×1 to 6×6, where the 6×6 kernel is equivalent to FC layers. DNNs with kernel size 1×1, 2×2, 3×3, 4×4, 5×5, and 6×6 achieved 0.53, 0.53, 0.56, 0.57, 0.57, and 0.57 ImageNet top-1 accuracies, respectively.

The presence of skip-connections was manipulated by replacing all convolutional layers in AlexNet with residual blocks of ResNet-18 (He et al., 2015). The model with skip-connections achieved 0.64 ImageNet top-1 accuracy. Similarly, the presence of branch-connection was manipulated by replacing all convolutional layers in AlexNet with inception blocks of Inception-v1 (Szegedy et al., 2014). The model with branch-connections achieved 0.58 ImageNet top-1 accuracy.

To manipulate the depth of a DNN, we inserted or removed additional convolutional layers into/from VGG-19 (Simonyan and Zisserman, 2014). Note that the additional convolutional layers were not included in the calculation of BH scores (i.e., the insertion of additional layers did not change the layer numbers). The default VGG-19 had depth of 19 (16 convolutional, two FC layers, and one category layers). The depth was reduced to eight by removing convolutional layers in each of five convolutional blocks except the representative layers. The depth was increased to 37 and 52 by inserting by inserting additional convolutional layers in each convolutional block: 0, 6, 4, 4, and 4 layers were inserted in the first to fifth convolutional blocks for depth of 37, and 5, 8, 8, 6, and 6 layers were inserted in the first to fifth convolutional blocks for depth of 52. These models achieved 0.58, 0.72, 0.63, and 0.68 ImageNet top-1 accuracies, respectively.

All DNNs were trained with an image category classification task on the ImageNet ILSVRC 2012 dataset. The batch size was 64 and the learning rate was 0.01, which was multiplied by 0.1 for every 20 epochs. The cost function was cross-entropy with L2 penalty. The coefficient of the L2 penalty term was 5 × 10^−4^. DNN weights were optimized by gradient descent with momentum (Qian, 1999) with a momentum term of 0.9. Dropout operations were inserted into every fully-connected layer except for the last layer. The dropout rate was set to 0.5. To prevent serious overfitting, we utilized early stopping based on the validation set.

### Statistical analysis

A permutation test was applied to examine whether the Spearman rank correlation coefficient between the BH score and image recognition performance is significantly deviated from zero (Figure 3). The tail probabilities of Spearman’s rho were computed using the Edgeworth series approximation (David et al., 1951).

### Data and Software Availability

The code to reproduce the results in this study is available at GitHub (https://github.com/KamitaniLab/BHscore). The fMRI data used in in the present study are available in public repositories; raw fMRI data are hosted at OpenfMRI (https://openneuro.org/datasets/ds001506) and preprocessed fMRI data are provided at figshare (https://doi.org/10.6084/m9.figshare.7033577). The stimulus images used in the fMRI experiment are available on request (https://forms.gle/ujvA34948Xg49jdn9). The unit activations of the DNNs as well as the decoded unit activations and the prediction accuracies are available at figshare (https://doi.org/10.6084/m9.figshare.12401168)

## Footnotes

a https://github.com/BVLC/caffe/wiki/Model-Zoo#models-used-by-the-vgg-team-in-ilsvrc-2014

b https://github.com/BVLC/caffe/wiki/Model-Zoo#models-from-the-bmvc-2014-paper-return-of-the-devil-in-the-details-delving-deep-into-convolutional-nets

c https://github.com/pytorch/vision/blob/master/torchvision/models/

d https://github.com/soeaver/caffe-model

e https://github.com/soeaver/caffe-model/tree/master/

f https://github.com/soeaver/caffe-model/tree/master/cls

g https://github.com/DeepScale/SqueezeNet

h https://github.com/shicai/DenseNet-Caffe

i https://github.com/tensorflow/models/tree/master/research/slim/nets

j https://github.com/tensorflow/models/tree/master/research/slim/nets

k https://github.com/dicarlolab/CORnet

